# Physical mechanism for the extremely rapid twinkling by chromatophores of Atherinidae

**DOI:** 10.1101/2025.10.25.684579

**Authors:** Masakazu Iwasaka

**Affiliations:** Hiroshima University

## Abstract

Lymphatics and secondary circulation systems of teleost exist in the skin layer where chromatophore cells coexist. This work reveals that in the skin of three species of Atherinidae, an extremely rapid flashing iridophore acts as fluid pressure sensor and visualizes micro-circulations in the layer. The fish has a capillary network propagating fluid pressure from the primal circulation systems. Complex of xanthophore, iridophore and micro-circulation network forms the twinkling spot that change light reflection intensity in 100 ms. The movement of light reflecting particles of iridophore, to which fluid pressure is applied, is the mechanism of twinkling skin tissue. The iridophores form a membrane with honeycomb compartments containing guanine platelets. Micro-circulation capillaries are embedded in the honeycomb membrane and provide driven forces for the light reflecting platelet movements. Two dimensional image of the micro-citculation appears on the twinkling patterns. The results suggest that real-time visualization of the micro-circulations can be achieved in the body of the Atherinidae. Making mimicry of the native skin device with the twinkling iridophores can contribute to developing the new artificial skin device for visualizing lymphatic micro-circulation system that can monitor an immune process in real time.

## 1 Introduction

In fish’s dermis and epidermis, the unique cellular system controlling colors is a group of chromatophore cells consisting of melanophore, xanthophore, iridophore, and so on (Fujii 1989). Among the pigment cells of fish, iridophore has a role to control light reflection intensity and adjust its colors (Denton 1970; Herring 2012). As a light reflecting particles of the iridophore, guanine platelets were investigated concerning ethological and optical aspects of aquatic animals (Jordan Partridge and Roberts 2012; Palmer, et al. 2017). Mechanisms responsible for the dynamic optical behaviors are attributed to the movements of the optically functional particles or protein fibers (Aihara, Kasukawa, Fujii and Oshima 2000). When a fish change its color of body surface, two mechanisms were proposed. One is the effect of aggregation and dispersion of light absorbing particles of melanophore, which is a kind of chromatophore for decreasing the light intensity inside body (Fujii 1989). The other is the possible inclination of light reflecting platelets (Denton 1970; Herring 2012; Iwasaka and Asada 2018).

Even though there were huge number of studies on fish chromatophore, no report is found on possible interaction of the chromatophores with lymphatic or second circulation. A previous study revealed that a skin tissue containing chromatophore of silverside fish (Atherinidae) exhibited dynamic light reflection phenomenon, twinkling of iridophores (Iwasaka 2021). This study verifies the hypothesis that the driven force for moving iridophore’s light reflecting platelets comes from the circulation systems, lymphatic or second circulation, of the silverside fish. Recent studies have shown that lymphatic vessels have organ-specific roles (Petrova and Koh 2020). Lymphatics and secondary vascular system of fish were discussed concerned with lymphatics of mammals in the past decades. It is remaining fundamental questions and insufficient understanding of the role of lymphatics in organ physiology. Revealing unknown features of lymphatic circulation system with micro-capillaries may stimulate biomedical engineering fields to bring new fluid dynamics device technology. From the viewpoint of biophysics, lymphatic system keeps unknown mechanism contributing to ingenious role of its micro-fluidic transport. Searching the new knowledge of lymphatics or secondary circulation was also carried out in the study of fish physiology (Petrova and Koh 2020; Sunyer 2013; Flajnik 2018). As a model for discovering new function of lymphatic systems, teleost fish’s immunity has high potential to provide new insight about the evolution of vertebrate immune system (Sunyer 2013). Understanding of vertebrate adaptive immune system was expanded by investigating fishes, cold-blooded vertebrates (Flajnik 2018). Previous studies on morphology and physiology of lymphatics of teleost fish brought a very important knowledge on fish lymphocytes which was useful for mammalian immune system study (Scapigliati, Fausto and Picchietti 2018). Lymphatics and secondary vascular systems of teleost fish were observed in the past centuries (Steffensen, Lomholt and Vogel 1986), and debates continued on analogies between fish secondary vascular system and mammalian lymphatic systems (Olson 2014; Rummer, Wang, Steffensen, Randall 1996). As a model organ, zebrafish was utilized for many of lymphatic investigations, because the fish was known to share major characteristics of lymphatics in mammalian (Jung, et al. 2017; Jensen, et al. 2009; Shin, et al. 2019).

If we are reminding about the coexistence of chromatophores with microcirculation systems in dermis, finding a possible concern of the microcirculation in functions of chromatophores should be beneficial to comparative immunology. In the past two decades, three species of fish were found to exhibit fast change of body surface color or shining intensity (Iwasaka 2021; Mäthger, Land, Siebeck and Marshall 2003; Goda 2017). This work reports that flathead silverside fish and cobalt silverside fish have a dynamic light reflecting spot, as well as hardyhead silverside fish (Iwasaka 2021). One of the features of twinkling light on body surface of the Atherinidae is the existence of strongly shining yellow region. This yellow light originates from capillaries forming a network around twinkling spot (TS).

## 2 Results

### 2.1 Light reflecting membrane on TS

On the dorsal trunk of silverside fish, circular spots consisted of chromatophores were discovered to cause rapid flash which was a light reflection intensity change. Typical image of this phenomenon is shown in Fig. 1a. The default color of the spot is green or blue and the size is around 100 µm. Remarkable point is that its flash speed is less than one second and the fish repeats the flash more than three times within one second. This repetitive flashing was not reported in the other two species (Mäthger, Land, Siebeck and Marshall 2003; Goda 2017). When the fish started this light flashing, time train pulses of strong yellow flash can be observed both by microscope camera and observer’s naked eye. In this study for the first time, it is found that the spot is covered by transparent membrane where iridophores align to form hexagonal pattern (Fig. 1b, Fig. S1). It was hard to recognize the existence of this transparent membrane when covering up melanophore. However, the honeycomb structure of conjugated iridophores, iridophore honeycomb (IrH), clearly appears in the gap of two melanophores, as shown in the picture of Fig. 1b and Fig. S2. The bridge part of IrH connects chromatophore spots and connected array of spots is formed, as shown in Fig. 1c. In addition, there are capillary-like structures distributing around the spots. A thick capillary connected two spots (Fig. 1c).

**FIGURE 1.**
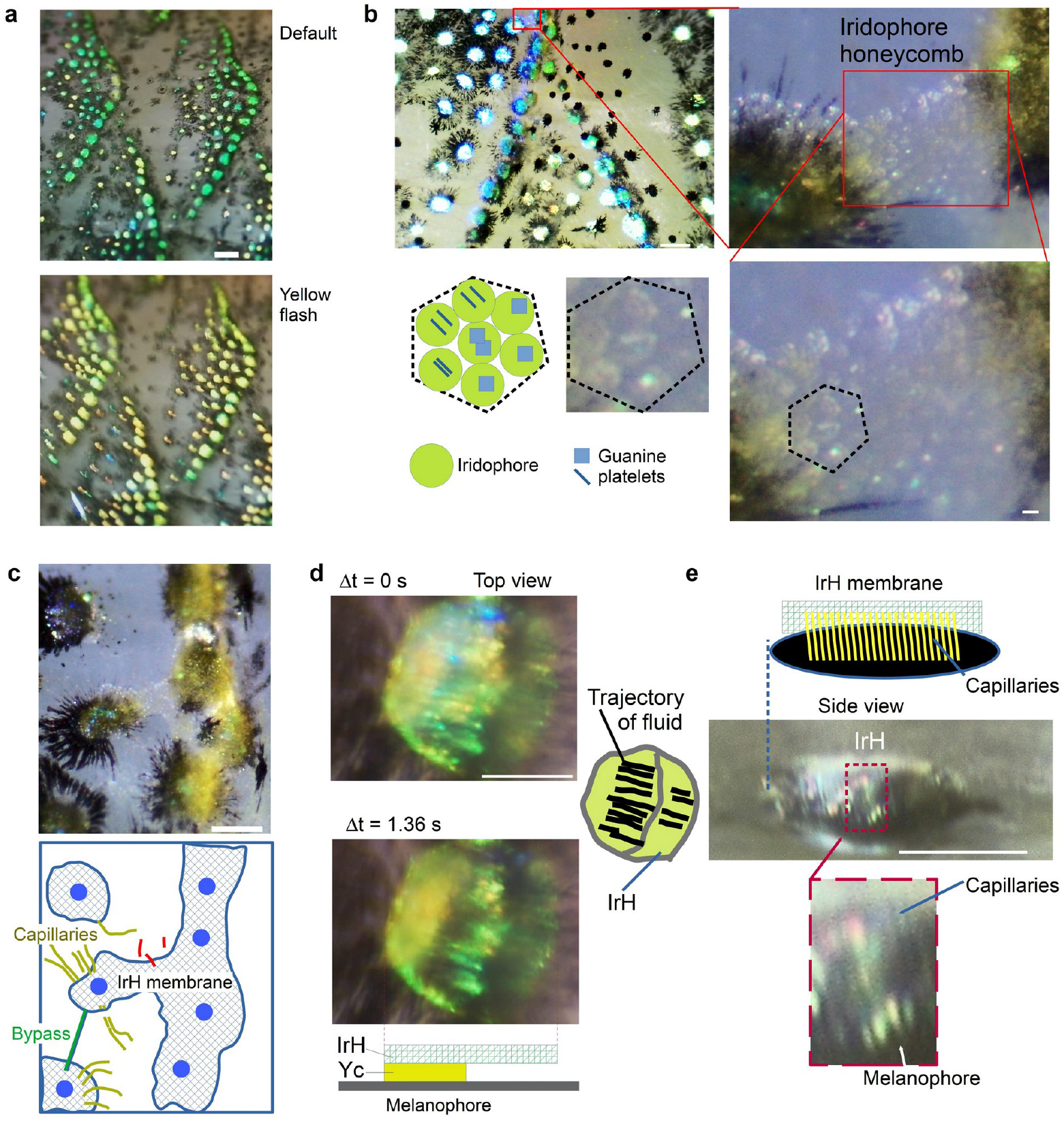
Membrane of iridophore honeycomb (IrH) covering the TSs of silverside fish’s dorsal trunk. (a) TSs changing color from blue/green to yellow. The spots exist in the edge of scale. (b) Microstructure of the spot-covering IrH. Inset with red line shows a IrH existing between two spots. In bottom panel, hexagonal inset with dashed line shows iridophores forming hexagonal pattern. Bright particles in the iridophore are guanine platelets. (c) IrH on multiple spots. Bottom panel illustrates IrH membrane (hatched area), position of spots (blue circle), and capillaries stretched around (ocherous line). A green line indicates a bypass-like capillary connecting two spots. Red line are blood vessels. (d) A time series of color changing spot where trajectory of fluid appeared (right hand illustration). Time span (Δt) of the two pictures is 1.36 s. Bottom illustration shows a speculated structure of the complex of IrH, Yc and melanophore. (e) Side view image of a spot. Capillaries adhering the spot side were observed. The picture was obtained in a tissue section. Species; (a)-(c), cobalt silverside, (d)-(e) hardyhead silverside. Scale bars, 100 µm.

Guanine platelets being packed inside the iridophore (Fig. 1b) produce structural color of blue. The color or light intensity changes provide information about the mechanical fluctuation in micro-scale region close to the platelets. From the view point of this mechanism, surface pattern difference between two photographs of Fig. 1d can be understood if mechanical forces such as fluid pressure is supplied for the IrH membrane. Fig. 1e shows the side view of *in vitro* sample’s spot. Beside the IrH covering black part of melanophore, bundle of capillaries whose diameter is several micrometers exist. Many parallel capillaries adhering to the side of spot are connecting to the IrH on top. An observation of the sectional plane of skin tissue indicated that yellow capillaries (Yc) distributed around and inside the spot (Fig. S3). Comparing images with different incident light angle identified guanine platelets existing within partially transparent spot as well as on the spot surface.

### 2.2 Micro-circulation capillaries around TS

Detailed analyses of the capillaries existing near IrH and spot were carried out (Fig. 2). On the body surface of silverside fish, there were areas containing yellow pigments around IrHs and melanophores. The area were seemed to overlap xanthophore from which the yellow pigment particles were supplied. In Fig. 2a, yellow and red line denote the capillary and blood vessel, respectively. Red arrows in pictures of Fig. 2b denote capillaries having blind-end. Bottom illustration and magnification of red box in Fig. 2c show capillaries overlapping on melanophore. In Fig. 2d, capillaries with radial arrangement exist separately from melanophore’s dendritic parts. Black arrow heads in magnification of red box of Fig. 2d denote reflecting particles remaining inside the capillary. Picture and its illustration in Fig. 2e show the yellow capillaries adhering to xanthophore.

**FIGURE 2.**
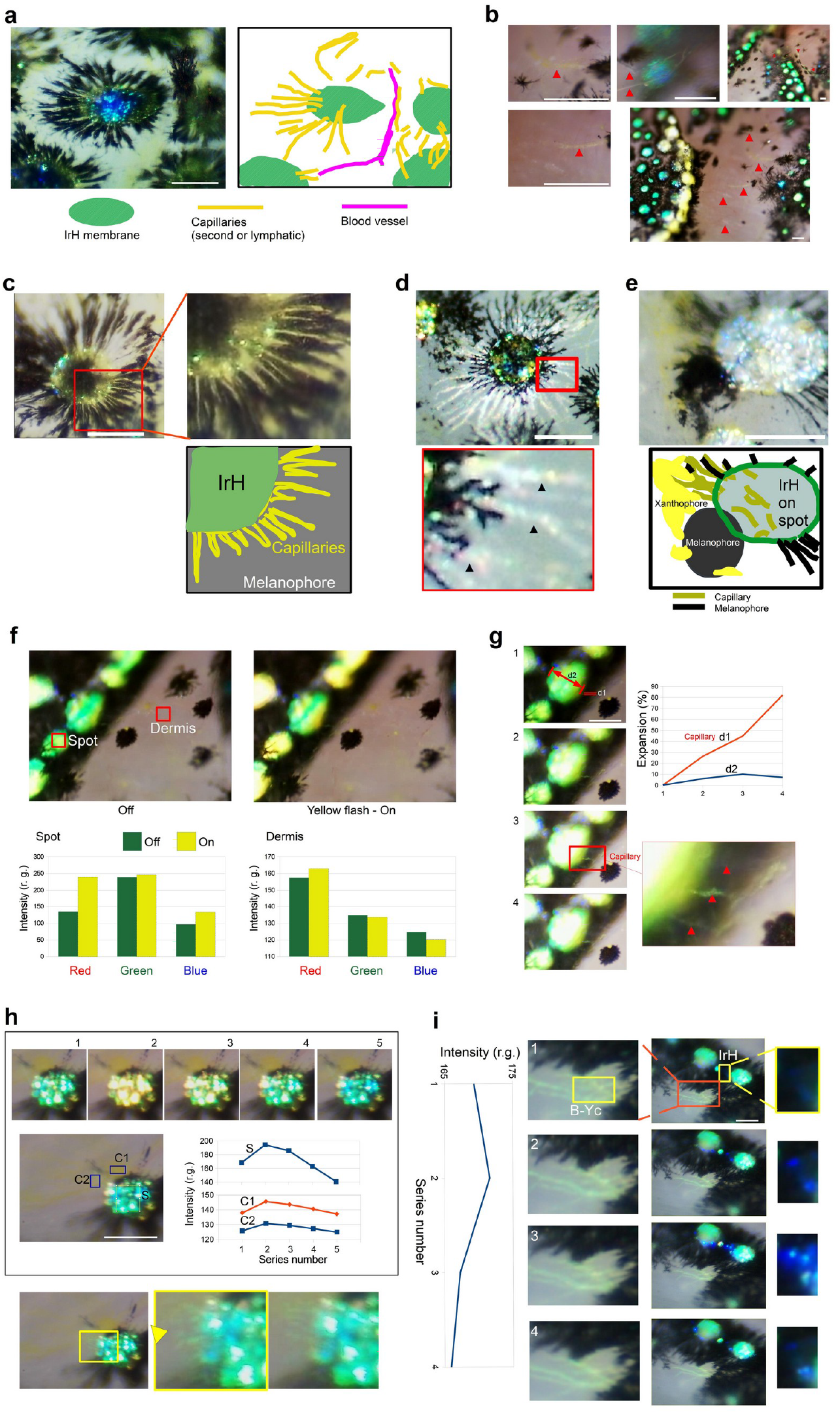
Structure and fluidic behaviors of circulation capillaries adhering to the TS of silverside fish. (a-b) Examples of appearance of capillaries surrounding and adhering to IrH on spots. (c-e) Distinction of the circulation capillaries from melanophore. (f) Correlation of blood circulation with the yellow flash in spot. Light intensity analysis of two regions (insets of left picture) is shown in bottom graphs. Intensity of RGB colors of a spot and dermis is obtained before and during yellow flash. (g) Effect of flash on the size of spot and capillary. Graph shows size change (%) of capillary width (d1) and spot diameter (d2). Red arrow heads of magnified inset (red box) denote capillaries. (h) Synchronization of light reflection intensity of spot and surrounding region of xanthophore and Yc. Five top images show transient process of twinkling. Time series of light intensity at three regions, on spot (S) and two surrounding regions (C1, C2), are shown. Fluid flow patterns appeared in IrH membrane layer (bottom). (i) Relationship between Yc’s fluid pressure and blue twinkling in circumferential part of spot. Scale bars, 100 µm.

In the macroscopic view of the trunk of the fish body, it was found that along the edge of scales there were tessellated pattern of the yellow pigmented area consisted of IrH and Yc (Fig. S4a). When observing surrounding part of a spot by optical microscope, xanthophores and blood vessels were found (Fig. S4b). In the gap of several spots, Yc were found as well as blood vessel, as shown in Fig. 2a. Also, yellow and blind-ended capillary were found out on epidermis of naked area distant from IrH-melanophore complexes (Fig. 2a-c). Usually the capillaries were yellow, however, white capillary was also found, as shown in Fig. 2d and Fig. S5b. Both yellow and white capillaries existed on the black dendritic parts of melanophore (Figs. 2c, d). The capillary existed separately from the dendritic melaophore. By applying noradrenaline to an euthanized body surface, the existence of the transparent capillaries was confirmed after its melanin aggregation completion (Figs. S6, S7). The location of the appeared transparent capillary was apart from territory of melanophore.

Two pictures in Fig. 2f are of without (left) and with (right) light flash on spots. Light intensities of three colors in spot and naked skin are shown in graphs for each case. The color of dermis which was observed through naked skin shifted to red when yellow flash took place in a spot (Fig. 2f). It suggests that fluid pressure of the capillary had correlation with blood circulation system. The Yc network should be fish’s secondary circulation or lymphatic.

In order to check the feature of fluid dynamics, relationships between spot and capillary was analyzed. Fig, 2g shows size change of fluid path when the pressure propagate from spot array to narrow capillary. Additional analyses (Fig. S8) support the evidence of the synchronization of spot twinkling and the sizes of connecting capillary when a fluid pressure pulse propagates. Correlation of light intensity fluctuation of a TS and Yc distributing around the spot was also obtained (Fig. 2h and Figs. S9-S13). There were cases where the reflected light fluctuation occurred in the same phase and different phases. Depending on the route of a fluid flow, the spatiotemporal pattern of light reflection was formed. The analyses of fluid pattern suggested that the capillary adhering to xanthophore connects to the overpass embedded in epidermis covering IrH (Figs. S13b,c).

Inset of Fig. 2d presents light reflecting particles are contained inside the white capillary. Certainly, a leakage of yellow pigments from xanthophore into capillary should occur. In addition, Fig, 2d indicates guanine particle leakages can enter the capillary from IrH to which the capillary connect. Possibly the capillaries act as both efferent and afferent ducts. The light reflecting particles were coexisted with yellow pigment particles and showed light blinking due to inner stream of the capillary (Fig. 2i and Figs. S14-15). An analysis of the reflecting particles under magnetic fields indicated that the particle oriented under magnetic fields in the same manner with guanine platelets (Fig. S16). Along the circumferential path of connecting spots (middle column pictures of Fig. 2i), blue flash lines are observed. This phenomenon frequently occur in cobalt silverside fish. When yellow pigments are localized in upstream, transparent fluid can cause distortion in IrH membrane and exhibit blue color which is peculiar to guanine platelet’s multiple light reflection.

Former two teleost species (Mäthger, Land, Siebeck and Marshall 2003; Goda 2017) showed rapid color changes within a second, but the process occurred once a second, and did not repeat within a second. However in the case of the Atherinidae, the circular iridophores housing guanine platelets exhibited repetitive change of color and light reflection intensity within a second. Its frequency of the change reached up to 3 Hz. So we need to consider a mechanism enabling the dynamic angle fluctuation of the guanine platelets. As well as iridophores of other species of teleost, the iridophore of silverside fish should have no ability to swing its guanine platelets repeatedly within a second by itself. A mechanical transducer should hide behind the IrH. The prime candidate is the discovered capillaries which can affect the inclination of platelets by transmitting fluid pressure to the iridophores of IrH. Flow pressure changes from the circulatory systems can be a driven force inducing a dynamic motion of the reflecting particles. The Yc network can induce enhancement of the yellow color on body.

### 2.3 Fluid pressure propagation visualization by chromatophore complex

It is apparent that the dynamic angle fluctuation of guanine platelets existing inside iridophore is the origin of rapid light reflection intensity changes. The obtained evidence of the optical behaviors of capillaries adhering to IrH indicates that the fluid of primal or secondary circulation is supplied to the capillaries, and a propagation wave of fluid pressure (Fig. 3a) acts as the driven force causing inclination of guanine platelets in IrH. Analyses of the capillaries using optical microscope shown in Fig. 1 and Fig. 2 bring the idea that an effective mechanical force can be transferred form the capillaries to iridophores. Based on this mechanism, schematic of distribution of capillaries in IrH-xanthophore-melanophore complex is produced (Fig. 3b).

**FIGURE 3.**
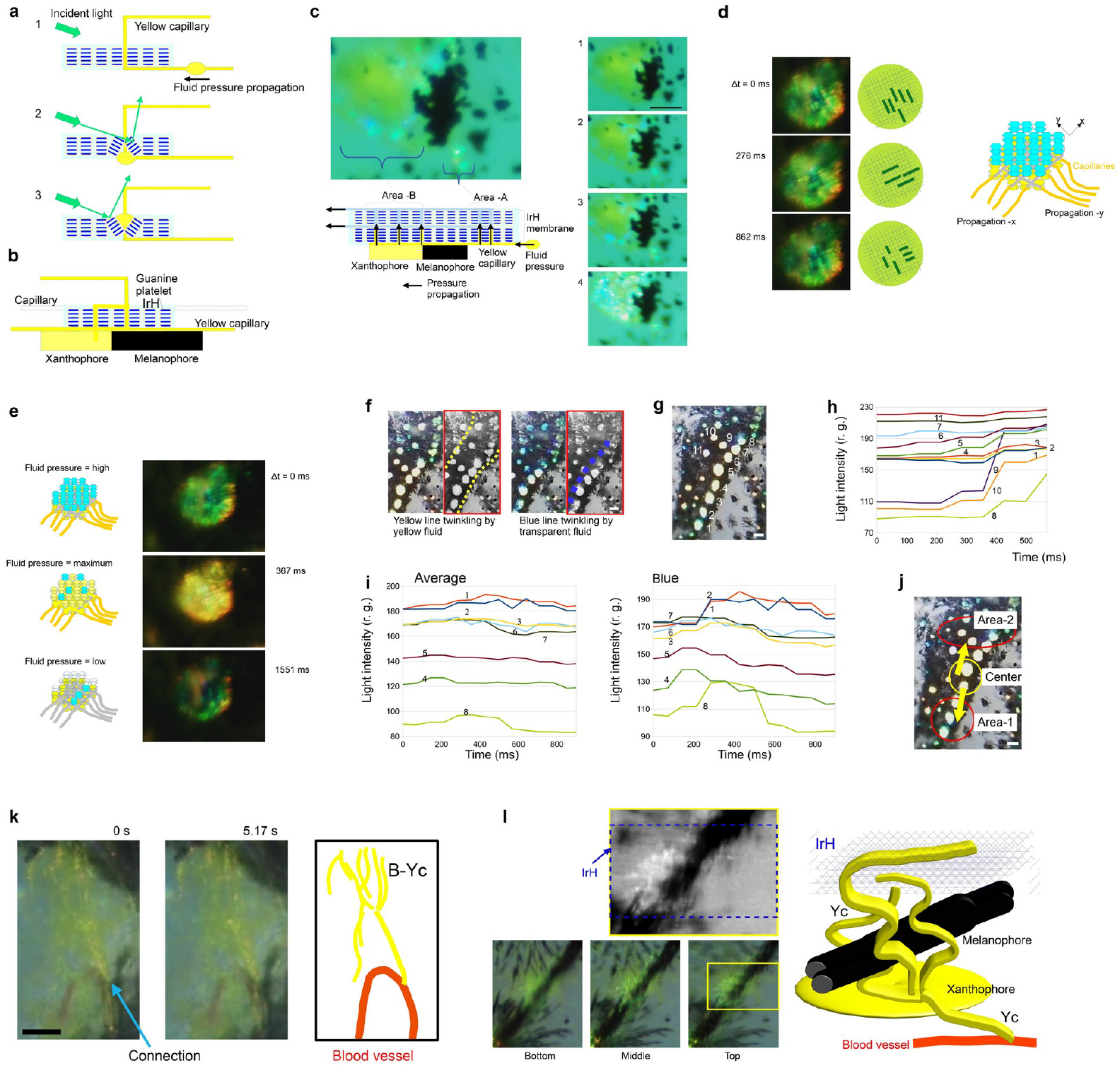
Mechanism of the visualization of secondary vascular fluid pressure propagation by IrH membrane. (a) Partial inclination of IrH causes strongly light reflecting area on the IrH surface. (b) Configuration of capillaries in the complex. (c) Evidence for melanophore-independent twinkling in IrH. Right panel shows time-series of four images of twinkling iridophore honeycomb on a xanthophore which is separated from melanopore. Left panel explains layered structure of the complex consisting of IrH, capillaries, xanthophore and melanophore. (d) Appearance of grid-shaped pattern on IrHs (left) when fluid pressures were supplied to capillaries along two directions (x and y). Illustrations show grid patterns in each picture (middle), schematic of the complex of IrH and bidirectional capillaries which can form the grid pattern (right). (e) Rapid yellow flash with a distinct contrast change on TS (right). Illustrations are indicating fluid pressure changes (left). (f-j) Fluid pressure propagation visualized by IrH on aligning spots. Scale bars, 100 µm. (k) Connection of a Yc to blood vessel. Two still images at different time show a slight change in yellow particles distribution. Right panel illustrates connecting part of Yc and blood vessel. (l) Melanophore-xanthophore complex. Scale bars, 100 µm.

In the observation of TS of silverside fishes, there was no evidence for the concern of melanophore because any part of melanophore was not moving when the rapid movement of light reflecting particles were appearing. In addition it was found that the twinkling was also provided by the circulation capillary on xanthophore in a melanophore-absent area, as shown in Fig. 3c (and Fig. S17). Aggregation of melanin particles of melanophore occurred in several minutes, however, blue color twinkling in isolated IrH occurred within 100 ms. Right photos of Fig. 3c presents guanine platelets inclining to generate blue color within IrH membrane on xanthophore. As shown in the conjectured model structure (Fig. 3c), layers of IrH containing capillaries lie on melanophore or xanthophore. Capillaries have regions with and without the yellow pigment particles of xanthophore. In this model, capillary in upper layer of IrH has less yellow particles. Pressure change in this transparent fluid can contribute to blue light reflection by guanine platelets.

A spot of the picture of Fig. 3d had a remarkable grid pattern which was a product of fluid pressure propagation in the capillaries in IrH of the spot. It is conjectured that the capillaries align along two crossed axes. The waves of propagation (-x and -y as illustrated in Fig. 3d) induce an inclination of guanine platelets of IrH. The occurrence of yellow flash can be understand in the manner shown in Fig. 3e. Three sketches in left panel of Fig. 3e describes IrH under a fluid pressure fluctuation. When the fluid pressure is maximum, yellow pigments in fluid are dispersed to the IrH area and contribute to enhance yellow light intensity. A spot in green color (default) and dark spot is generated under high and low pressure, respectively.

Fig. 3f shows appearance of yellow line (left picture) and blue line (right picture) of TSs on edge of scale. Black-and-white picture in red box has dashed lines which indicates the area where yellow or blue line appears. Figs. 3h-j show analyzed two time-series of reflected light flash dynamics in the spots array of numbered spots shown in Fig. 3g. Based on the RGB-averaged light intensity and blue only data (Fig. 3i), the fluid path is conjectured (Fig. 3j).

In the example shown in Fig. 3h, spots No. 4-7 had a gradual increase of light intensity in the beginning, then spots No. 8-10 and No, 1-3 became brilliant. Another example (Fig. 3i) has the same tendency when the data were evaluated on average of RGB values, and the blue light intensity presented a clear evidence supporting this evidence. These data of time-series of individual spot’s brightness indicates that the pressure propagates from spot No. 4-5 to both No.1 and No. 8. Provably a circulation fluid pressure came out from dermis beneath center area of the picture in Fig. 3j, and propagated to two directions along the spot array.

The results indicate that the fluid pressure concerns a primal controlling mechanism of time-sequential twinkling of spots forming into a line on edge of the fish’s scale. In a zebrafish’s study, streams of lymphatic in epidermis were found to form tessellated lymphoid network pattern matching for the outline of fish scales (Robertson, et al. 2023). We can explain that the fluid pressure in the capillary adhering to TS came from second vascular and lymphatic vessels.

As the dynamic light reflection of spot in milli second order is driven by changes in capillary’s pressure and perfusion, the dynamic signal contain information about immunological activity of fish. Sometimes yellow flashing spots appeared at the same time over two to three scale covering area. In this occasion, we can make additional hypothesis that fast increase of the fluid pressure in blood volume propagated into capillaries connecting to spots.

## 3 Discussion

Recent research on vascular physiology was opened to discuss on specialized blood vessels forming under the developmental process of lymphatic systems of a mammal (Das, et al. 2022). Existence of lymphatic and second vasculature systems in fishes is promoting the discussion on diversity of tissue microcirculation systems in animals evolution (Panara, Varaliová, Wilting, Koltowska and Jeltsch 2024). In the present study of a flathead silverside fish, a new feature of fish microcirculation systems was discovered (Fig. 3k). The photographs in Fig. 3k show a Yc directly connecting to a blood vessel. Yc and xanthophore closing to a blood vessel were frequently observed (Fig. S18-S20). Among the data, the Yc shown in Fig. 3g has the apparent connection to the blood vessel. In the video playback of the data (Supporting movie S5), the blood flow dynamics correlated with the fluctuation of particles inside the Yc. The particles fluctuation was caused by the fluid motion that was provided due to a leaking of blood flow waves via the connection. The yellow stains in surrounding area of spots is a part of a xanthophore. Because the capillary Yc is embedded in IrH, guanine particles leaking from broken iridophore can enter into the capillary. Owing to the complex of Yc and IrH containing guanine platelets, the spot twinkling can involve physiological information about vasculature systems.

The vascular pressure is affected by contraction and relaxation of vessel, nitric oxide and flow volume (Gashev 2008; Gasheva, Zawieja and Gashev 2006). The IrH membrane over dermis of silverside fish is capable to elicit a visual detection of lymphatic vascular functions such as lymphatic pumping mechanism, shear stress interaction in lymphatic endothelial cells and fluid oscillation with a higher frequency of phasic contraction (Gashev 2008; Gasheva, Zawieja and Gashev 2006; Kunert, Baish, Liao, Padera and Munn 2015). Frequency of phasic contractions of lymphatic vessels of rat was reported to be approximately 0.2 Hz in average (Gasheva, Zawieja and Gashev 2006). A simulation model with calcium and nitric oxide level change showed the presence of a higher frequency of phasic contraction (Kunert, Baish, Liao, Padera and Munn 2015). The silverside fish’s twinkling spot covers frequencies of 0.1 Hz to 3 Hz (Iwasaka 2021). A nerve innervation of the second/lymphatic vasculature contraction should exits.

Fig. 3l suggests the formation of TS proceeds with the interaction between melanophore and the capillaries of lymphatic or secondary circulation. Three bottom-left images in Fig. 3l are of different layers (top, middle, bottom), which are obtained by changing the focusing. Left-top image of Fig. 3l (yellow box of top layer) under contrast adjustment shows IrH. It is known that cellular interactions between melanophore and xanthophore is the key process for the stripe formation in fish skin (Yamaguchi, Yoshimoto and Kondo 2007; Nakamasu, Takahashi, Kanbe and Kondo 2009; Mahalwar, Walderich, Singh, Nüsslein-Volhard 2014). IrH is already formed in the epidermis of Fig. 3l, however, capillaries adhering to xanthophore are coiling around the bundle-shaped melanophore as illustrated. The structure is the incomplete or destroyed chromatophore complex. The evidence of the chromatophore complex visualizing fluid pressure propagation by guanine platelets can become a new model for lymphatic imaging technique. In conclusion, the basic principle of rapid light reflection switching by the fish guanine platelet has a feasibility of enhancing visualization of micro circulations in skin of silverside fish. Particularly, the light reflecting tiny platelets can act as an optical transducer for physiological signals from second circulation and lymphatic whose inner fluid is linked with the primal vasculature system. Observing iridescent dynamics of the fish may enable us to accelerate learning another story of immune system evolution from fish. The fishes have dermal/epidermal photonic device for the display of micro circulation systems on living body.

## 4 Materials and Methods

### 4.1 Sample

Specimens of three species of Atherinidae were captured in Okinawa, Japan, by using a net and in Fukuoka and Hiroshima, Japan by fishing. The specimens were kept in aquariums (23–28°C), and up to 72 specimens were observed for this study. The height (and weight) of *Atherinomorus lacunosus* (hardyhead silverside), *Hypoatherina valenciennei* (sumatran silverside), and *Hypoatherina tsurugae* (cobalt silverside) were 108.4 mm ± 10.2 mm (13.9 g ± 2.6 g), 91.7 mm ± 11.6 mm (9.1 g ± 3.1 g) and 113.0 mm ± 9.5 mm (16.0 g ± 3.9 g), respectively. The specimens were selected without distinction of sex. The study was approved by the institutional Animal Care and Use Committee (approval number: 2F19–2, Hiroshima University) and carried out in accordance with all national guidelines including the guidelines for proper conduct of animal experiments (Science Council of Japan).

### 4.2 Observation of twinkling spots in fish’s body surface

*In vivo* observation of light reflection tissue was carried out in the same manner with the reported study (Iwasaka 2021) where an aquarium accommodating anesthetized fish was utilized. In the monitoring aquarium, the fish under anesthesia control was laid as previously described (Iwasaka 2021). The fish’s anaesthetic condition was controlled by 2-phenoxyethanol (0.1%, Fujifilm Wako Pure Chemical Co., Japan) and tricaine (50 to 100 ng/ml, Tokyo Chemical Industry, Japan).

By utilizing microscope lens (NAVITAR 2.0 × 1-51473, Navitar, Rochester, USA), CMOS camera (HOZAN L-835, HOZAN, Osaka, Japan) and a LED light source (LA-HDF158A, Hayashi Repic Co. Ltd, Tokyo, Japan), a time series images (video images and static images) of body surface of anesthetized fish was obtained.

A white LED light (LA-HDF158A, Hayashi Repic Co. Ltd, Tokyo, Japan) was used as a light source. The light illumination was provided from a light guide whose inner diameter was approximately 8 mm. The center axis of the incident light was directed towards the back of the fish, as shown in Fig. S21. Directions of incident light on top view and side view are described. The illumination directed from anterior to posterior. On the top view, vector of the incident light was in parallel to antero-posterior axis (Fig. S21). As shown in the side view, incident angle of light irradiated to water surface covering top of dorsal trunk was 47 to 56 degrees. Inclination angle of the fish’s trunk surface versus the water level was within ±5 degrees. The microscope lens focused in length of 20 mm – 30 mm, and image consisted of the reflected light from the specimen was monitored.

Throughout the text, following abbreviations are used for referring to specific structures found in the study.

- twinkling spot: TS
- yellow capillary: Yc
- iridophore honeycomb: IrH
- brilliant yellow capillary: B-Yc

### 4.3 Observation of tissue *in vitro*

Chromatophores and capillaries in the epidermis and scale was also observed in silverside fish under euthanasia with overdose of tricaine. A thin slice of tissue was set between glass slide and cover glass, and a cut surface was observed to find a spot showing side views by the same optical microscope mentioned above.

In addition, a neurotransmitter stimulation of dorsal trunk was carried out to induce melanin aggregation by using noradrenaline (1 mg/ml, Alfresa Fharma, Japan). The treatment was carried out to inspect the structure of skin when melanin particles cause aggregation increasing the tissue transparency.

### 4.4 Image capture and analysis

For the movie recordings of optical microscope images, a video capture software (Xploview, VIXEN, Saitama, Japan) was used. The white balance parameter was fixed during the individual movie recording. The flame rate per second (fps) of the recording was 12 fps to 30 fps. Still images involving a region of interest (ROI) were extracted from the movie, and mage analyses were carried out using Image J (NIH) software. When evaluating the reflected light intensity correlating with the microcirculatory dynamics, gray values measured in the software were used as the intensity. The intensity relative to gray is described as r.g. in the main text and Supplementary Materials. Average or individual gray value of three colors (RGB) provided by the software was obtained.

### 4.5 Analysis of brilliant particles of xanthophore under magnetic fields

An *in vivo* observation of brilliant particles in a capillary (B-Yc) was carried out on an anesthetized fish. The plastic-box type of aquarium accommodating anesthetized fish was set in the gap of magnetic poles of an electromagnet. The gap length was 70 mm and the electromagnet generated magnetic fields of up to 500 mT (Iwasaka, et al. 2013; Iwasaka and Mizukawa 2013). Light illumination was applied in parallel to the body axis of fish at the same incident angle with the cases of the other *In vivo* observations. The direction of magnetic field was perpendicular to the body axis and Light illumination. This combination of light illumination, magnetic field and vision provides a light reflection enhancement when guanine platelets in skin tissue orient in parallel to the magnetic fields, as reported in the previous works(Iwasaka, et al. 2013; Iwasaka and Mizukawa 2013). By using the same optical microscope system mentioned above, video movies of the brilliant particles of xanthophore on dorsal trunk were recorded before, during and after the magnetic field exposure at 500 mT.

## Data availability

Source data (Images and videos) and dataset of the presented figures are available at Zenodo after publication.

All other data supporting the findings are available within the paper and its Supporting Information.

## Acknowledgments

Author appreciates the support from JST-CREST Advanced core technology for creation and practical utilization of innovative properties and functions based upon optics and photonics (grant no. JPMJCR16N1).

## Conflict of Interest Statement

Author declares that we have no competing interests.

## Notes

### Competing Interest Statement

The authors have declared no competing interest.

