## Supporting_Information for "Physical mechanism for the extremely rapid twinkling by chromatophores of Atherinidae"

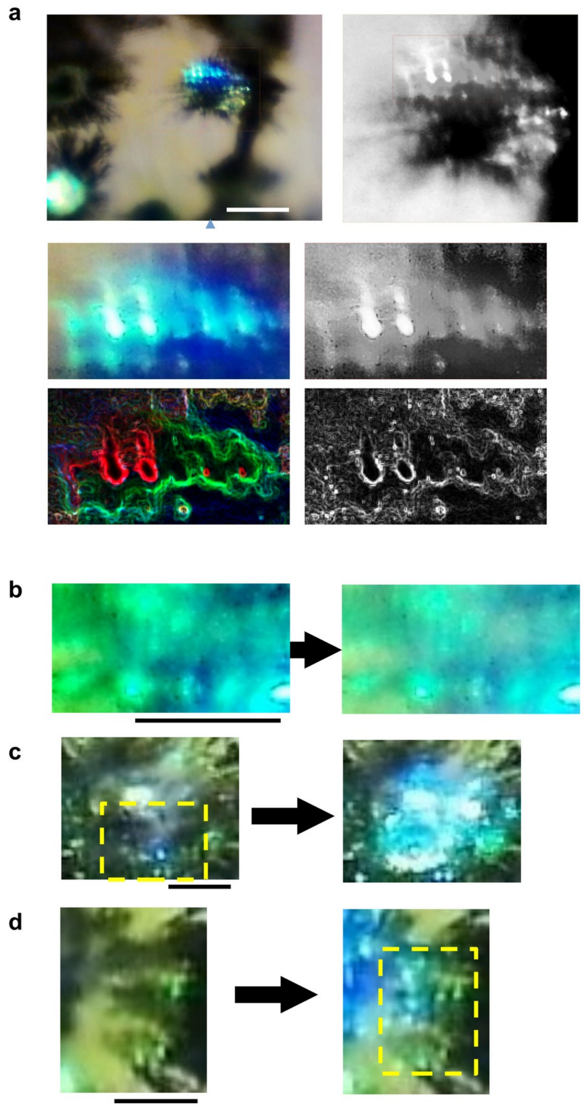


**FIGURE S1** | IrH membrane covering top of a twinkling spot of cobalt silverside. (a) Insets of the original picture were shown in black-white, magnified image and edge-enhanced images. (b to d) Texture changes of IrH when twinkling occurred. Broken yellow boxes show the place where iridophore can be observed as blue points. Scale bars, a, 100 µm, b to d, 50 µm.

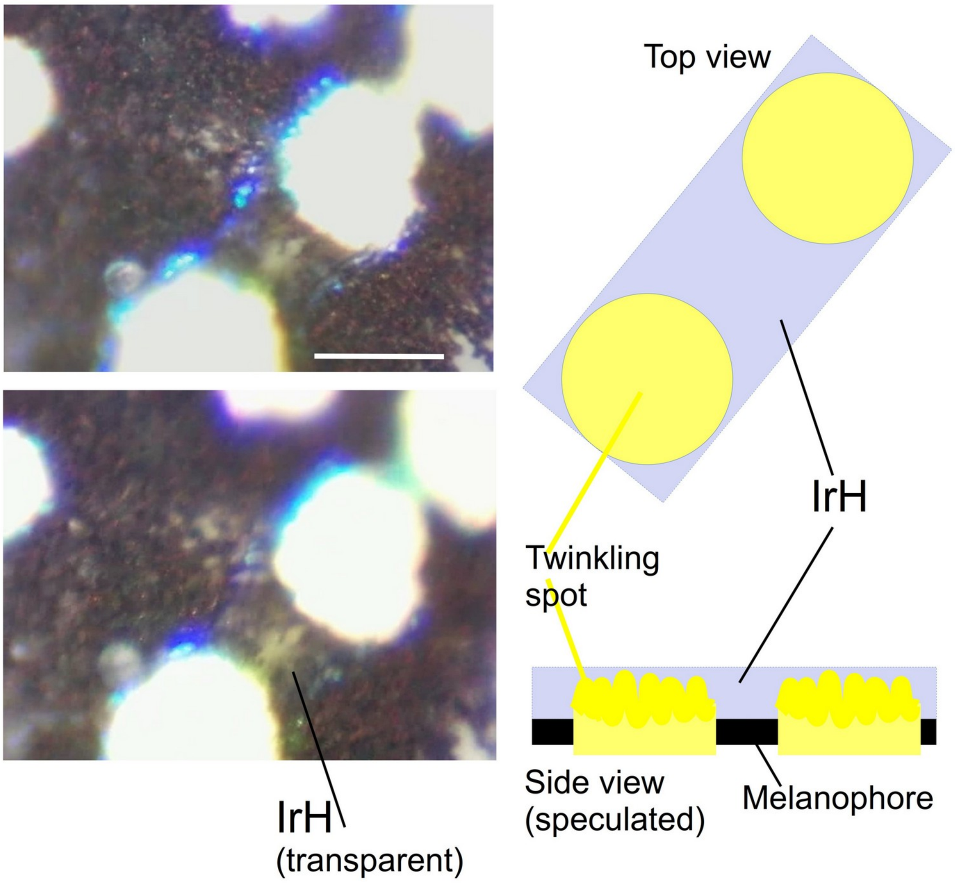


**FIGURE S2 |** IrH forming a bridge which locates over two spots of cobalt silverside. The transparent membrane of iridophores has a boundary with blue light reflection. In bottom layer of the complex, yellow pigments of YC can be seen. Right illustration shows speculated structure of the complex of melanophore (black), twinkling spot (yellow) and IrH. Scale bar, 100 µm.

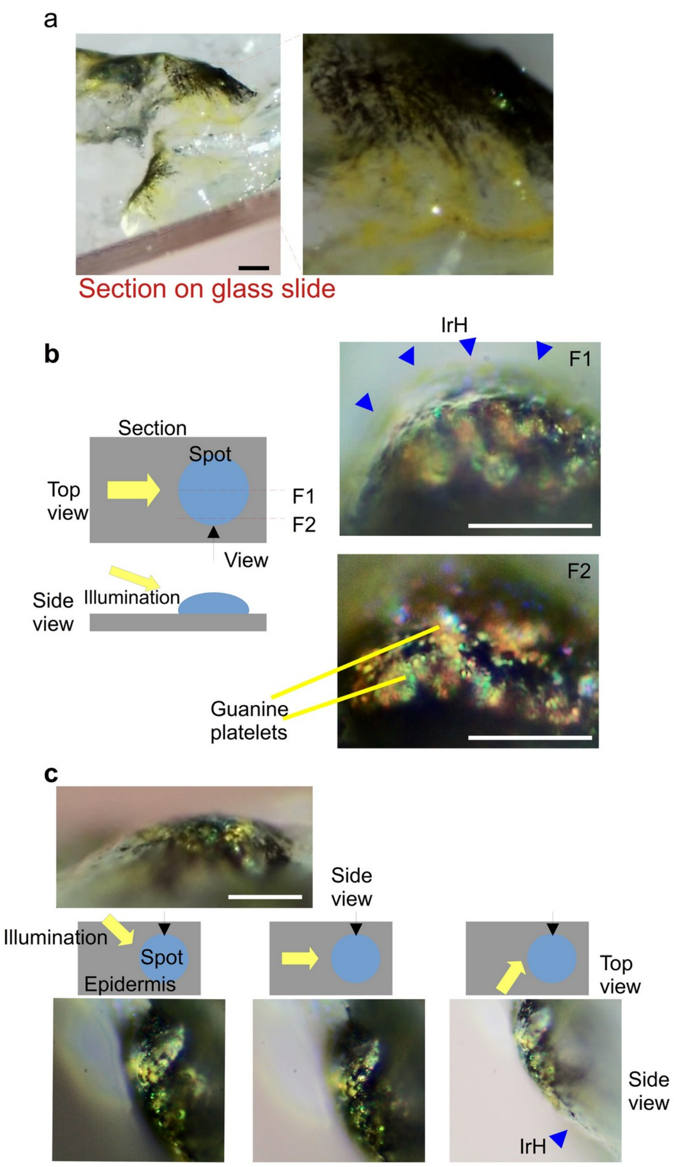


**FIGURE S3 |** Internal structure of twinkling spot observed in a section of hardyhead silverside. (a) Capillaries with yellow pigment are observed around melanophore. (b) Two side-view pictures under side illumination. Focusing were set at plane F1 and F2 of the left panel for top and bottom picture, respectively. Blue arrow head denotes IrH. Yellow lines denotes guanine platelets. (c) Still images of a spot observed from a side when illumination angles were set in thee cases. Scale bars, a, 200 mm, b and c, 50 mm.

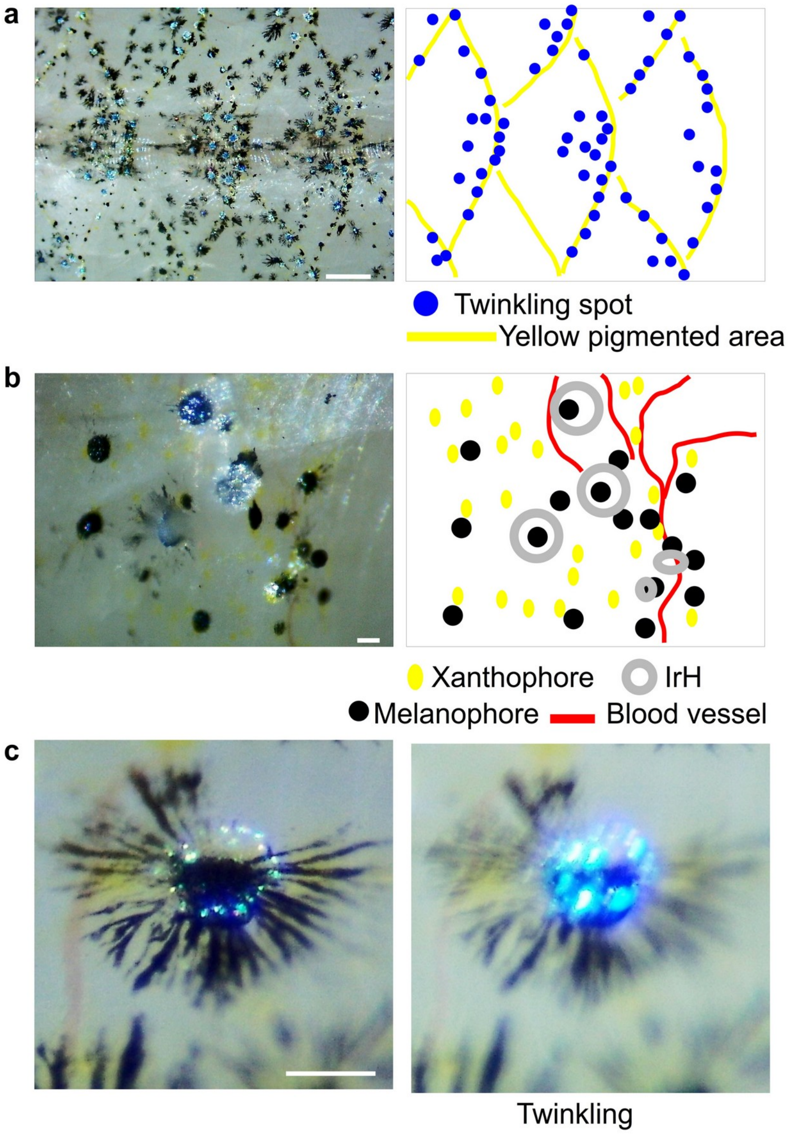


**FIGURE S4** | Arrangements of twinkling spots, blood vessels and chromatophores in dorsal trunk of flathead silverside fish. (a) Tessellated pattern of the IrH with Yc (yellow pigmented area) distributing along the edge of scales. Twinkling spots are also aligning along the edges, (b) Region of isolated xanthophores scattered around melanophores. Blood vessels and IrH membrane can be observed. (c) A melanophore accompanying IrH, xanthophore and blood vessel. Scale bars, a, 1 mm, b and c, 100 mm.

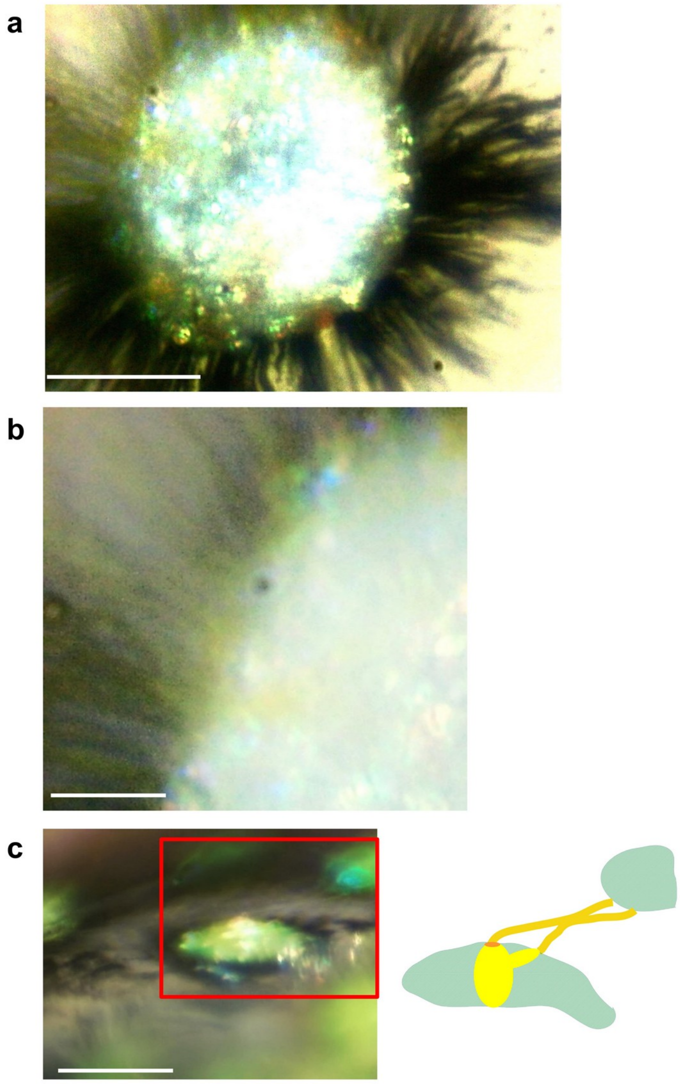


**FIGURE S5** | Capillaries existing on the surface of spot and epidermis of cobalt silverside fish. (a) Capillaries are radiating and looks like being embedded in IrH. (b) Yellow and white capillaries can be observed in the top layer of a spot. (c) Capillaries form a terminal of yellow pigments on top of spot. In the red box, there are two white capillaries connecting two spots, as illustrated in right drawing. Scale bars, a, 50 µm, b, 10 µm and c, 100 µm.

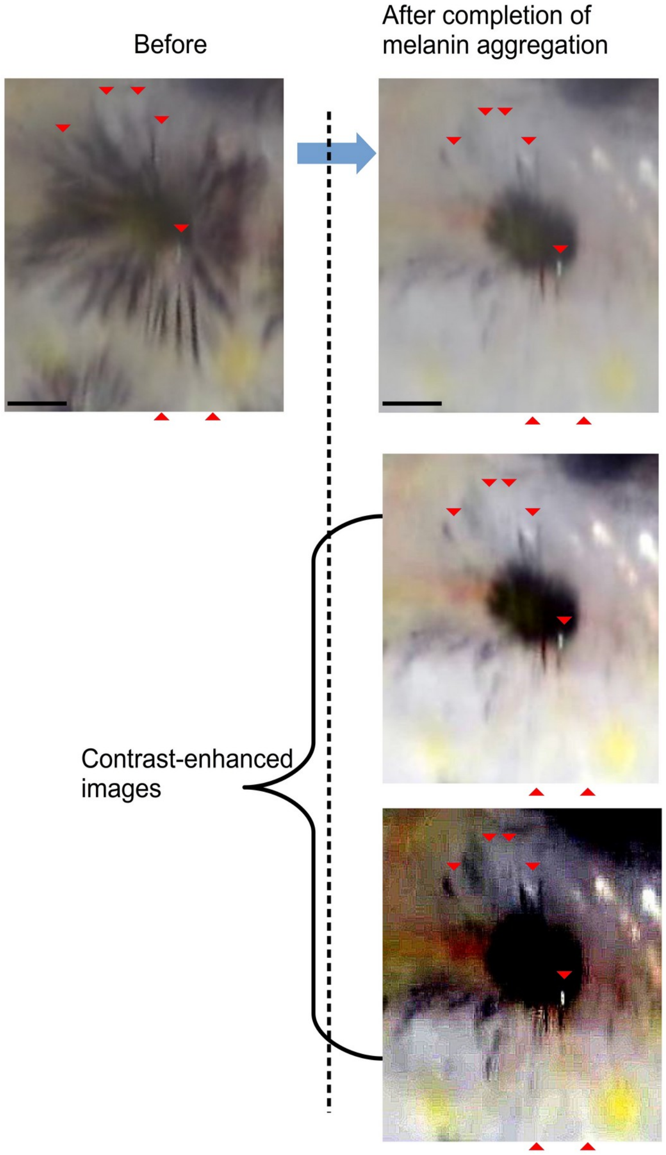


**FIGURE S6 |** Distinction of capillaries belonging to Yc with dendric part of melanophore.  Left picture shows a melanophore before the stimulation by norepinephrin. Some white capillaries are existing between the black radiating parts of melanophore. Right pictures are of after melanin aggregation. Bottom pictures show contrast-enhanced images. Independent from the melanin aggregation, the white capillaries were kept in the same locations. Scale bar, 100 µm.

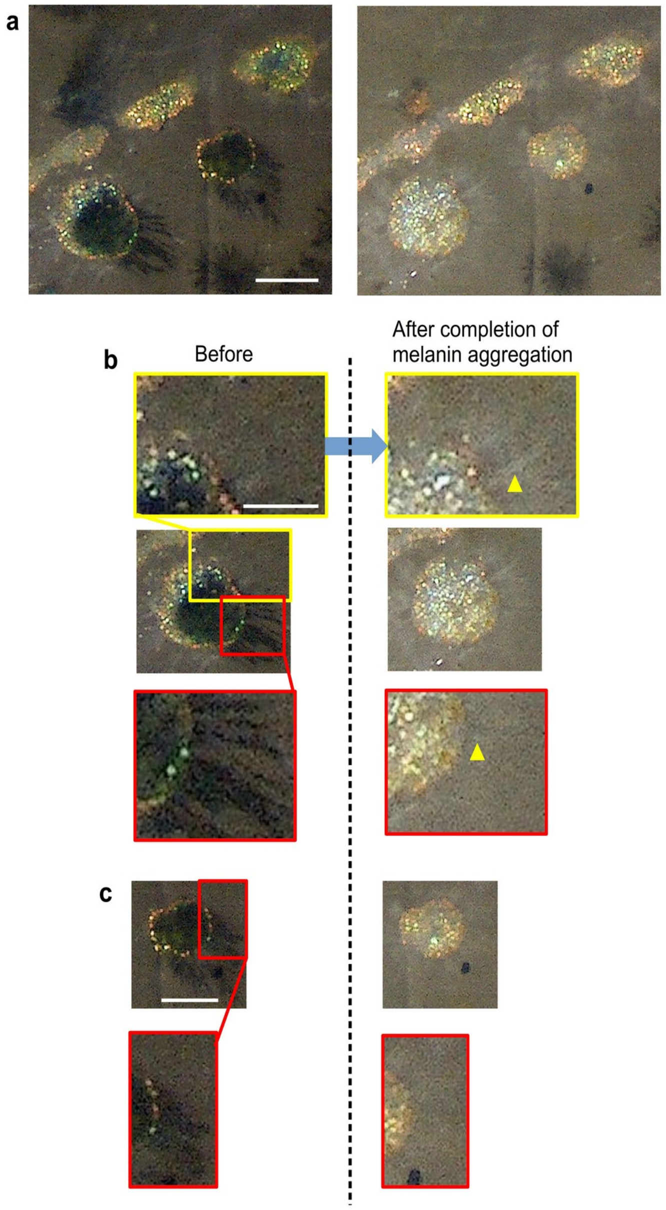


**FIGURE S7 |** Location of capillaries belonging to Yc and the dendritic part of melanophore. Pictures of spots before (left) and after (right) the melanin aggregation. (a) Still images in macroscopic view. (b, c) Searching the location of white capillaries whose locations are different from those of melanophore. Yellow arrowhead denote the example of white capillary independent from the melanophore. Scale bars, a, and c, 100 mm, b, 50 mm.

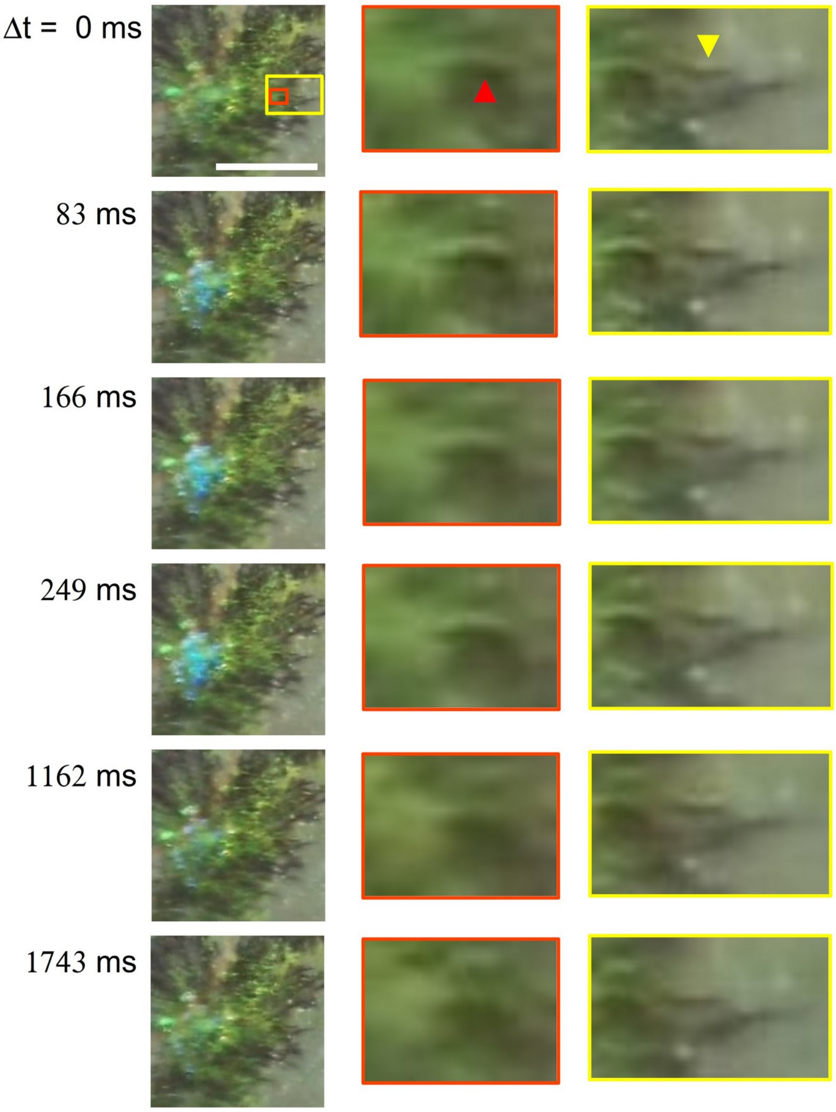


**FIGURE S8** | Slight swelling of capillary at the occasion of spot twinkling. Left panel show a twinkling spot showing blue light reflection in 83 ms to 249 ms. Its insets are magnified in middle and right panels. Arrow heads denote swelling capillaries. Scale bar, 100 μm.

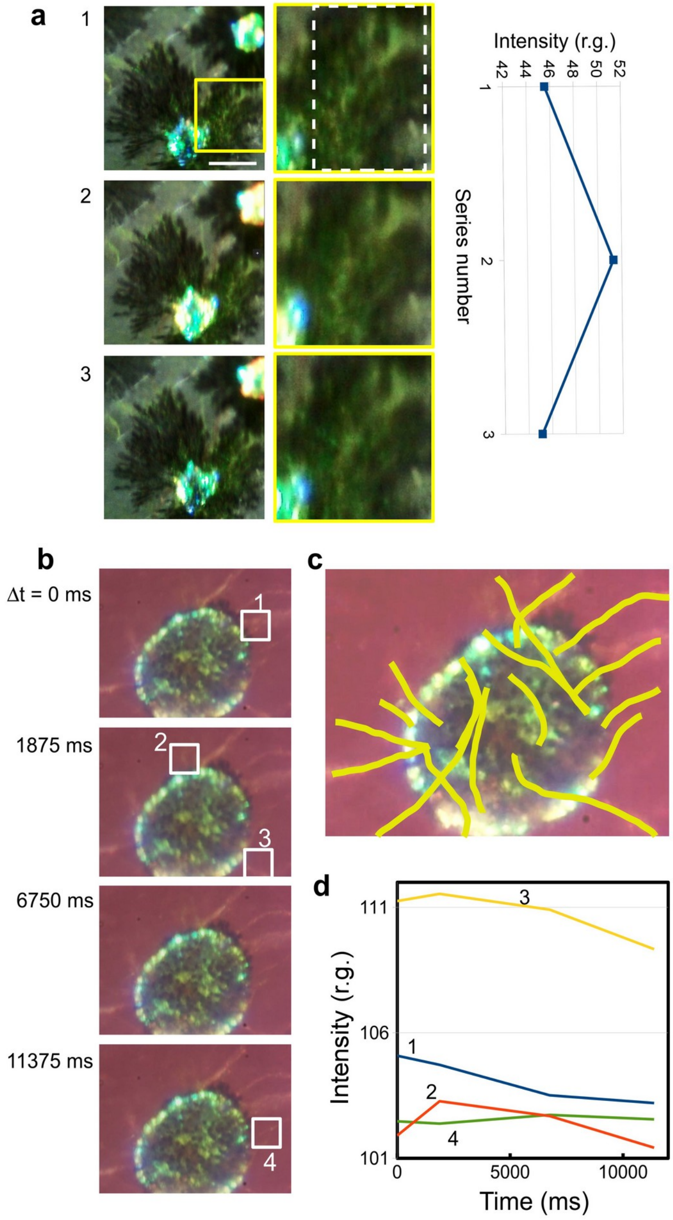


**FIGURE S9 |** Analysis of correlation of light reflection intensity of spot and capillary part. (a) Example of 'in phase' occurrence. Scale bar, 100 μm. (b) Fluctuation of light reflection in capillaries of a tissue under euthanasia. Yellow trace lines on picture (c) represent the capillaries found in picture of (b). Light intensity analysis on four part of capillary (inset-1 to 4) is shown in a time-series graph (d).

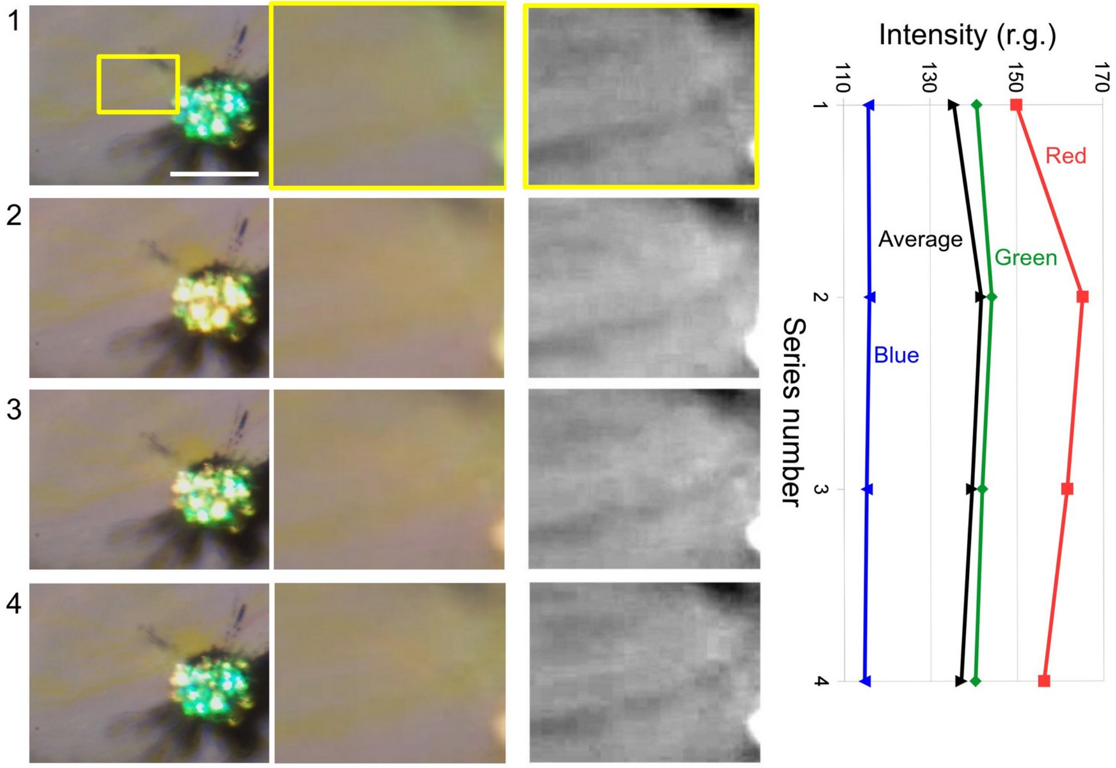


**FIGURE S10 |** Synchronization of yellow flash of spot and YC's behaviors (radiance and sweelling). Second column is the inset of top picture of the first column. Its RGB and averaged intensity relative to gray is shown in right panel. Third column is the black and white images converted from the second column, and provides information about the canal's swelling. Scale bar, 100 µm.

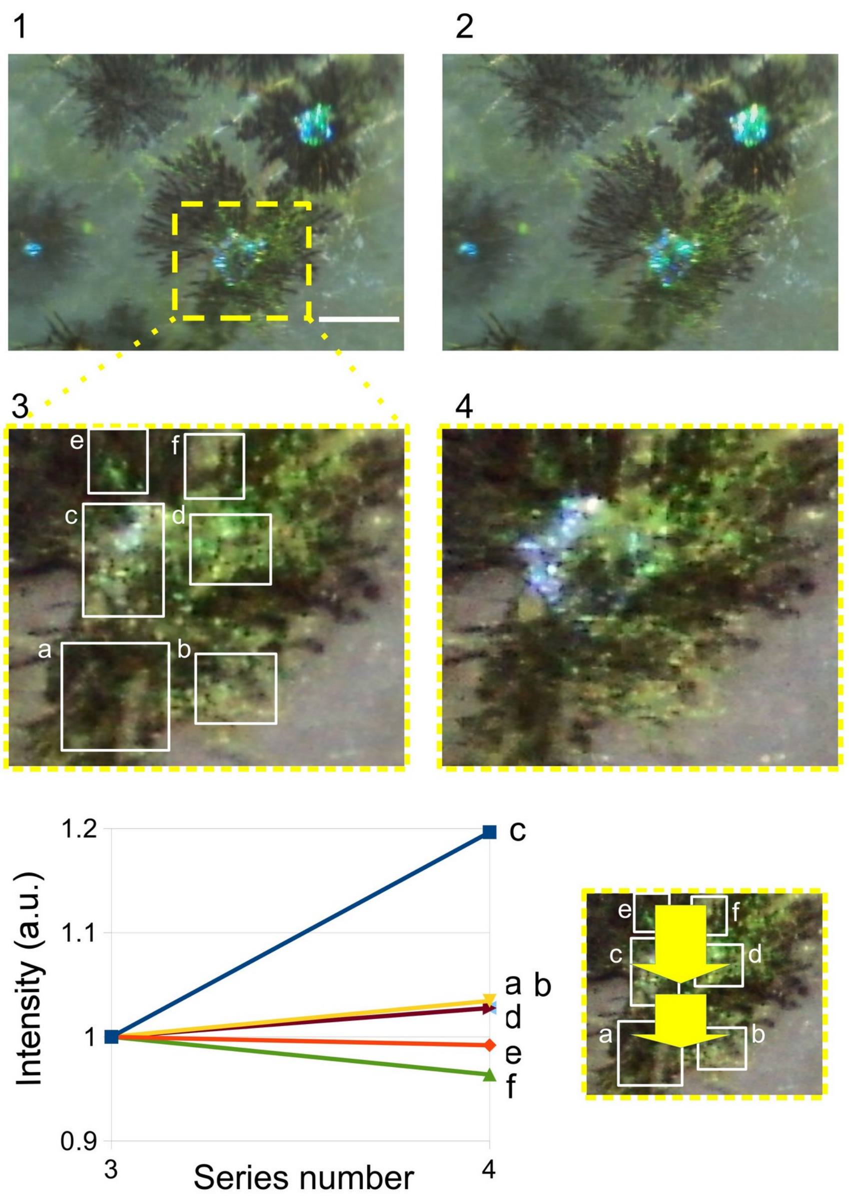


**FIGURE S11 |** Analysis of light reflection intensity distribution relating to fluid pressure propagation in and around a spot. Pictures 1 to 4 show a time series of twinkling spot. Bottom left panel shows normalized intensities (arbitrary unit) of light reflection in six areas. Area c and d belong to the spot. In bottom right panel, yellow arrows indicate a speculated path of pressure propagation. Scale bar, 100 µm.

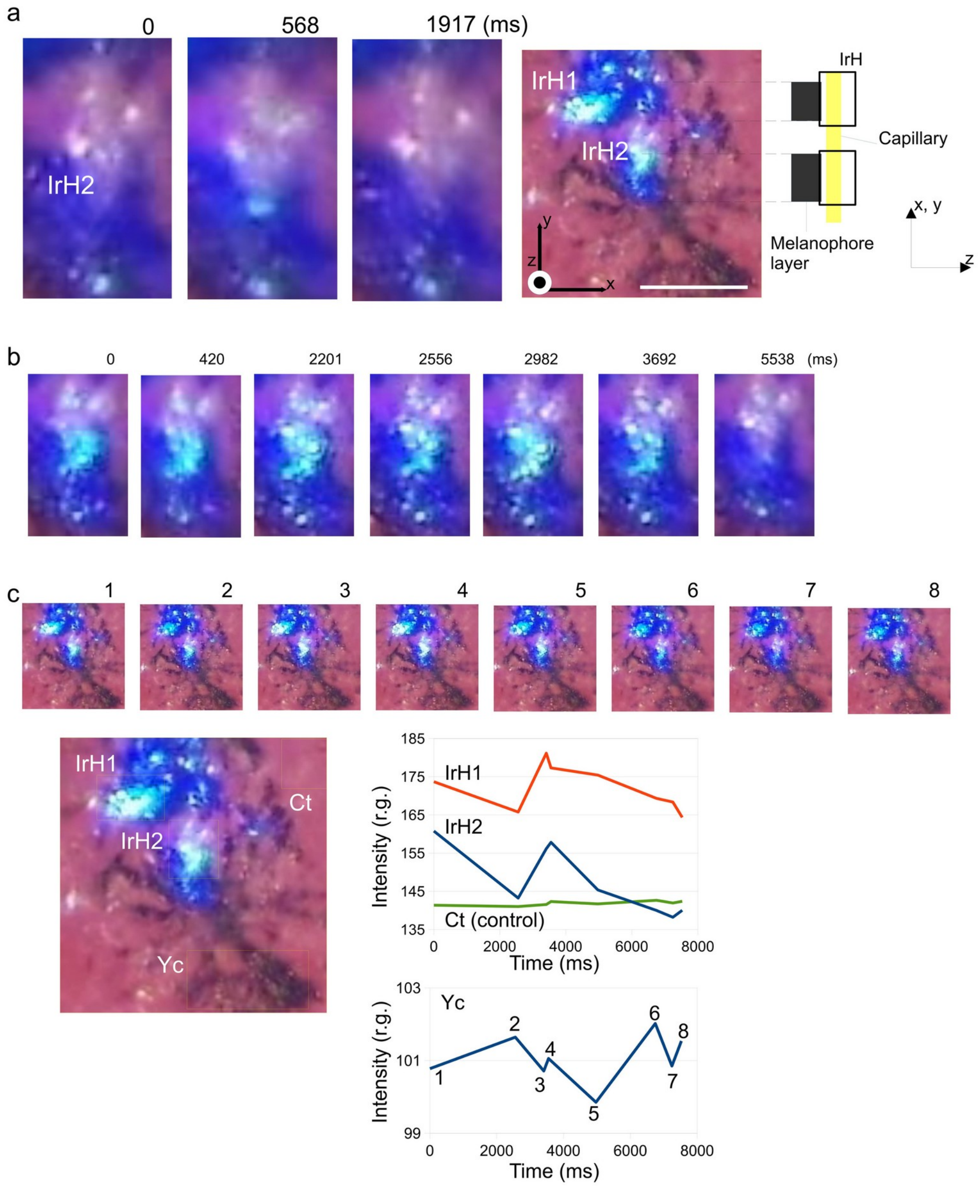


**FIGURE. S12 |** Example of weak correlation of light reflection between two IrHs and a Yc. (a) IrH2, one of two parts of IrH shows slight change in light reflection. Right panel shows speculated structure of the chromatophore complex. (b) IrH2 showing light reflection going down. (c) Analysis of the reflection intensity of two IrHs and surrounding two areas (control area (Ct) and capillary (Yc) in bottom of the picture). The Yc area had weak correlation with the IrH areas, because the Yc was out of pressure propagation path. Scale bar, 100 μm.

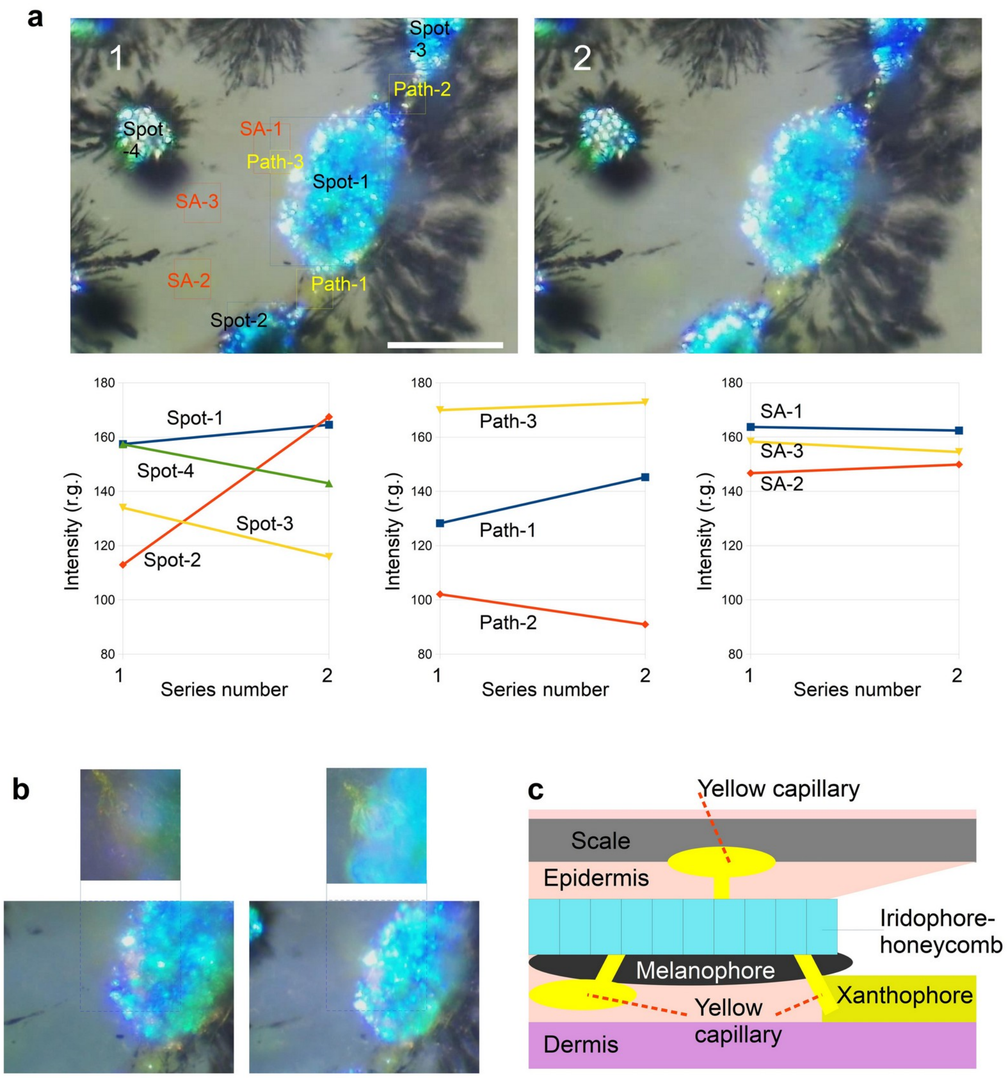


**FIGURE. S13 |** Analysis of light reflection intensity distribution in he field of multiple fluid pressure paths. (a) Light reflection intensity in four spots, three paths and three surrounding areas (SAs) are analyzed. The intensity of spot-1 & 2 increased while those of spot-3 & 4 decreased. This suggests fluid pressure propagated from top to bottom of the picture. Intensity increases of path-1 & path-3 also suggests this speculation. In addition, comparison of intensities of three SAs indicates the fluid pressure shifted to the bottom. (b) Two pictures show the existence of a capillary having a route to upper layer. (c) Stacked layers of melanophore, IrH, Yc and YC in epidermis beneath the scale. Scale bars, 100 mm.

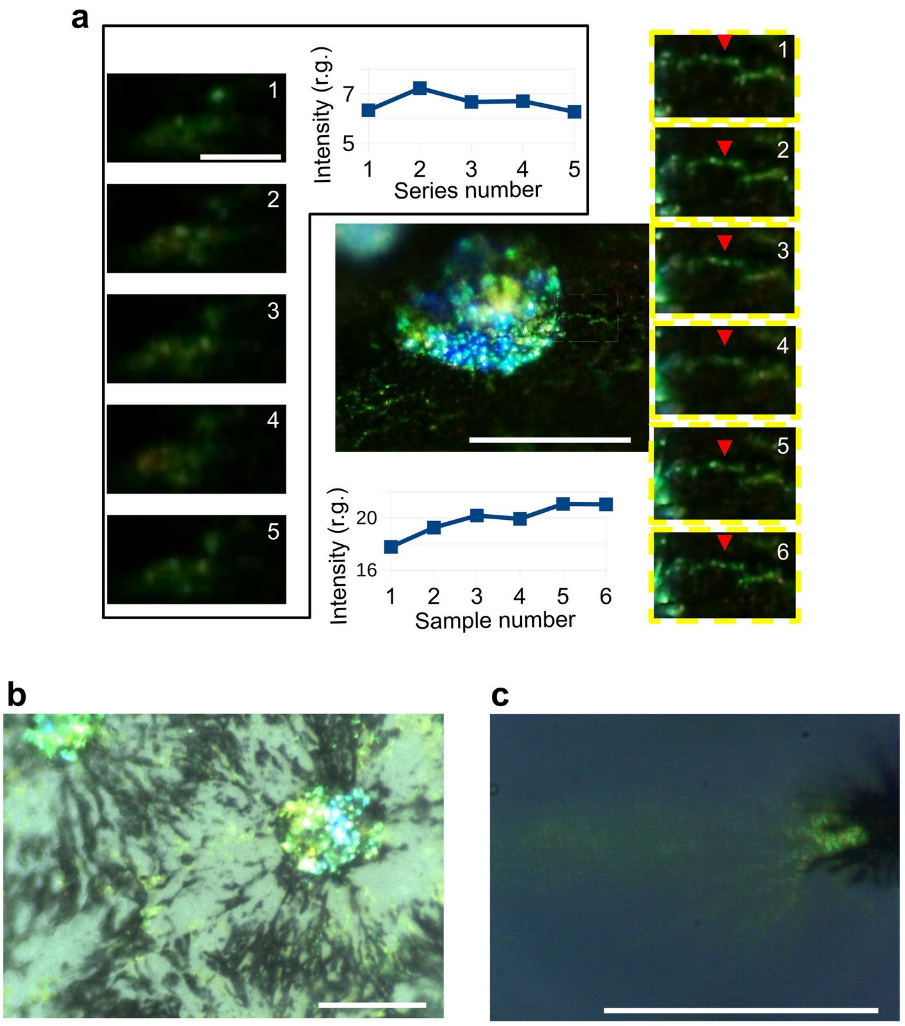


**FIGURE S14 |** Analysis of the movement of brilliant particles existing inside a yellow capillary of *Atherinomorus lacunosus*. (a) Time series of  captured images of the brilliant yellow capillary (B-Yc) showing light reflection fluctuation. (b and c) Two examples of B-Yc. Scale bars, a (left), 10 μm, a (right), b and c 100 μm.

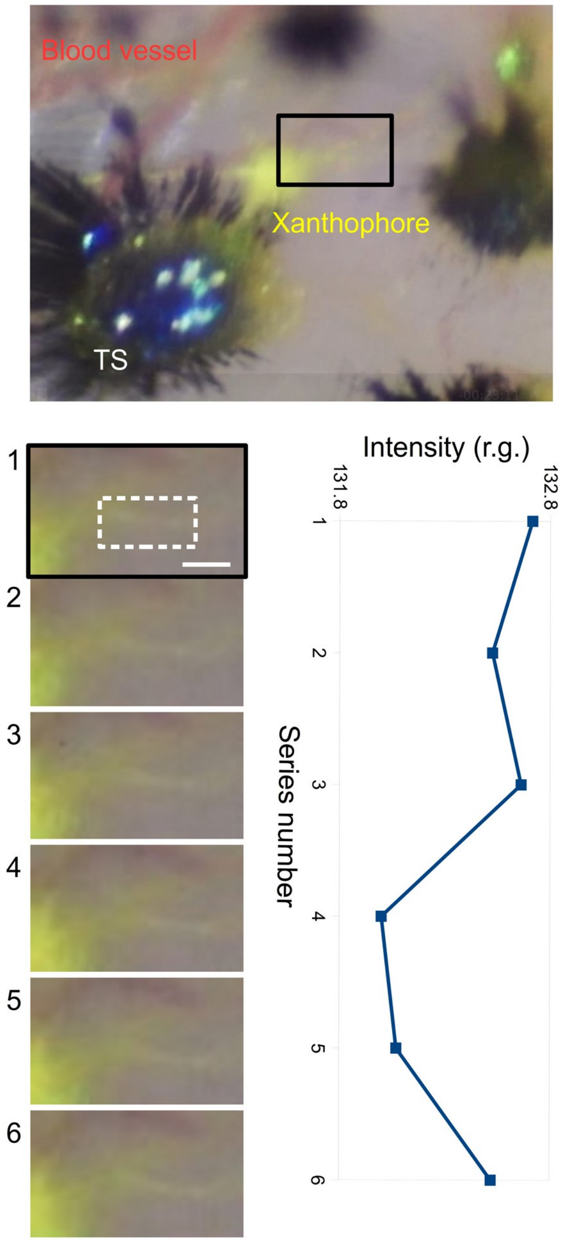


**FIGURE S15 |** Analysis of the B-Yc dynamics. In the center of the top picture, a xanthophore keeping its dendric shape is elongating its B-Yc. Bottom panel shows time series of the B-Yc of xanthophore. The movements of particles is detected as light reflection fluctuation (bottom right panel). Scale bar, 10 µm.

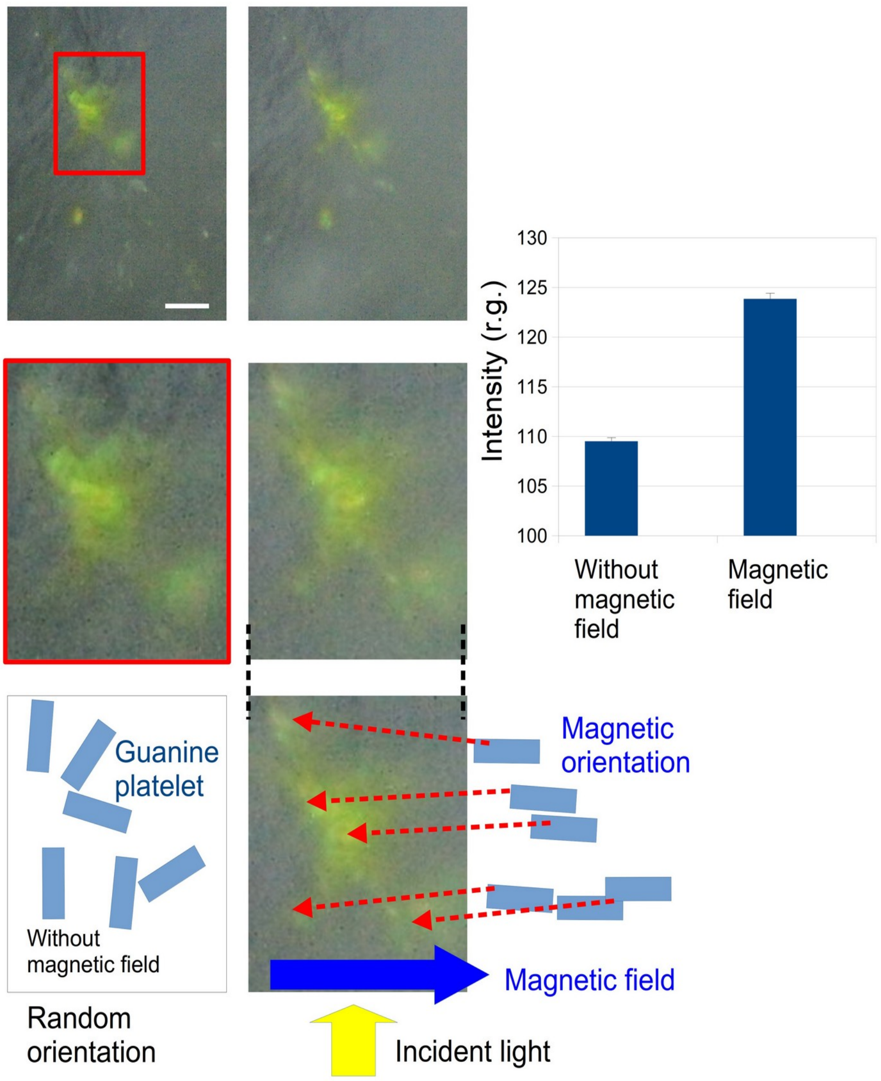


**FIGURE S16 |** Magnetic response of the reflecting particles contained in a brilliant Yc (B-Yc). Picture in left column show B-Yc in the absence of magnetic field exposure. Right column show pictures of B-Yc with a magnetic field at 500 mT. Detailed explanation of the magnetic field exposure is described in the method section. The region of red box inset is magnified and its light reflection intensity is analyzed for both cases, with and without the magnetic field. Tight graph shows intensity relative to gray of the regions of interest, which suggests the light reflection by the B-Yc was enhanced under magnetic field. Bottom drawing interprets the occurrence of magnetic orientation of guanine platelets existing in the Yc. The combination of magnetic field, incident light and oriented platelets provides the enhancement of light reflection. Scale bar, 10 µm.

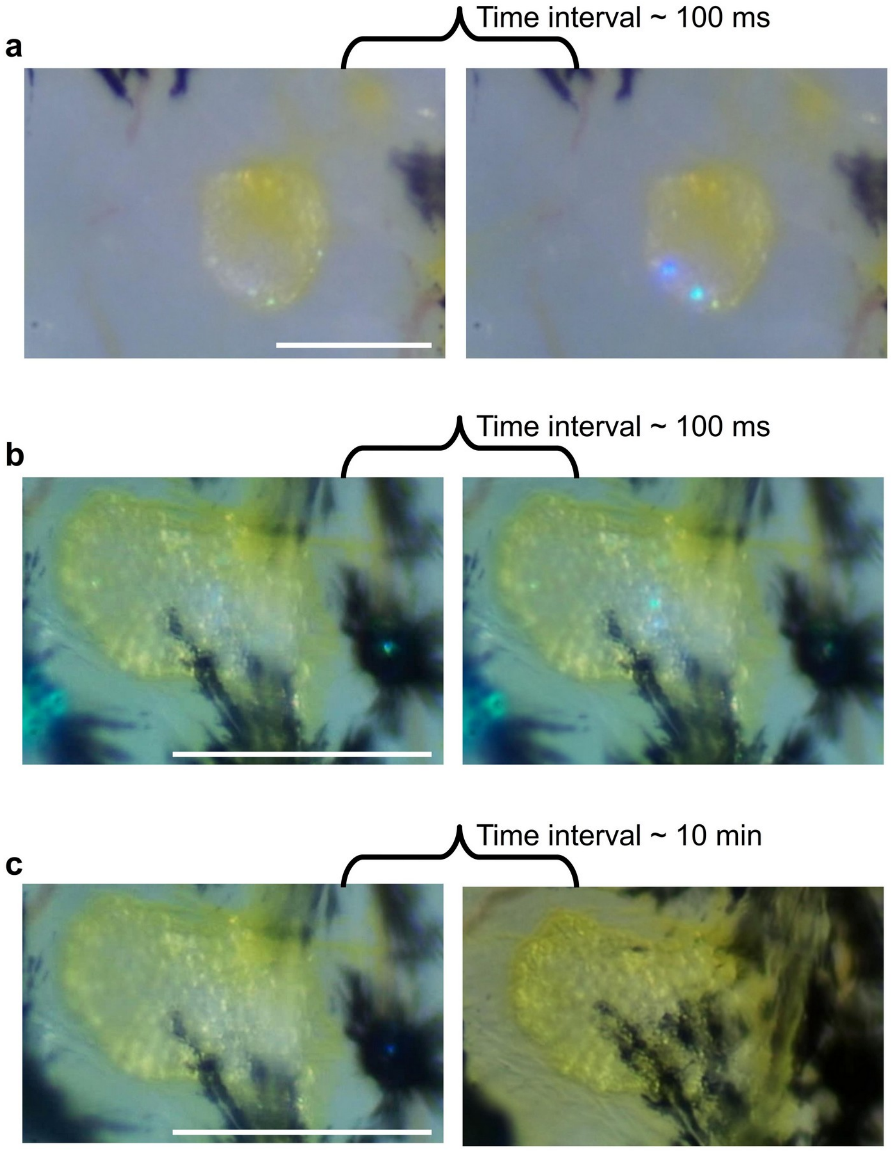


**FIGURE S17** | No concern of melanophore's aggregation dynamics in the twinkling of IrH. (a) Rapid twinkling in IrH membrane which is apart from melanophore. (b) IrH existing on edges of two melanophores. Black dendritic parts of melanophores are situated beneath the IrH. Pictures show the blue/green twinkling occurred in 100 ms. (c) No distinct effect of melanin dispersion on the light reflection of IrH. Some black dendritic parts of melanophore are elongating in 10 min, but no twinkling occurred. Scale bars, 100 mm.

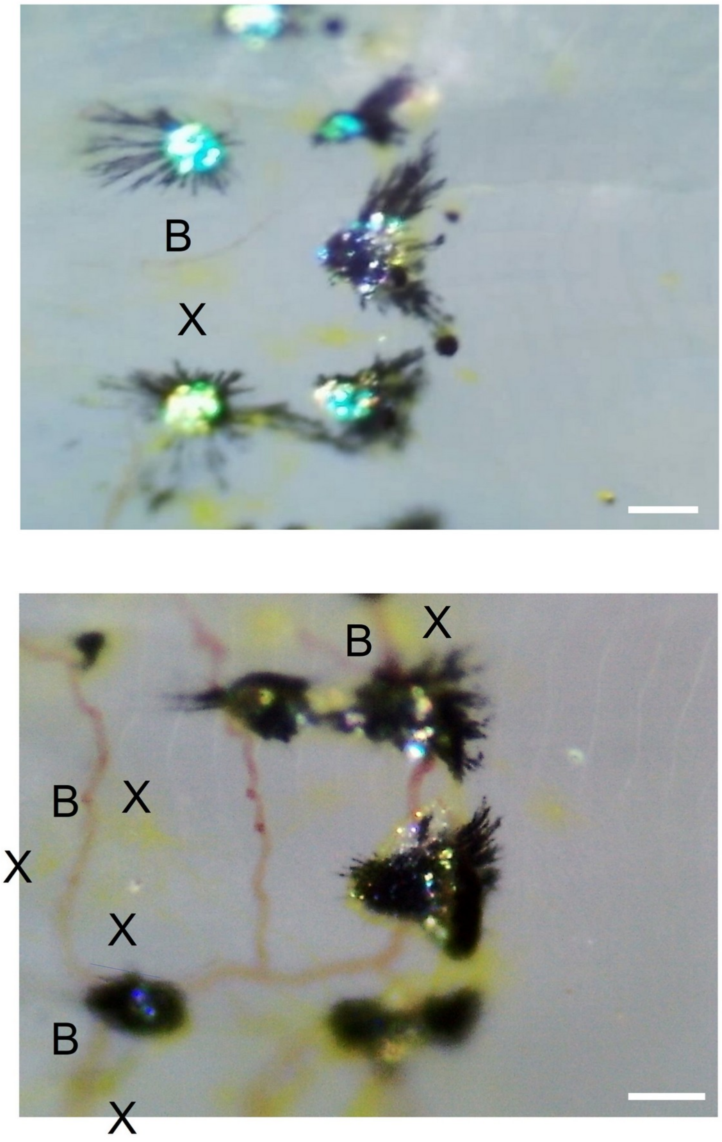


**FIGURE S18 |** Arrangements of xanthophores (X), blood vessel (B) and twinkling spots in epidermis. Top picture shows a xanthophore (or yellow canal) coupled with a blood vessel with faint red. Bottom picture is the case with a group of xanthophores and blood vessel. There can be seen a tendency of the xanthophore being attracted to a blood vessel. Scale bars, 100 μm.

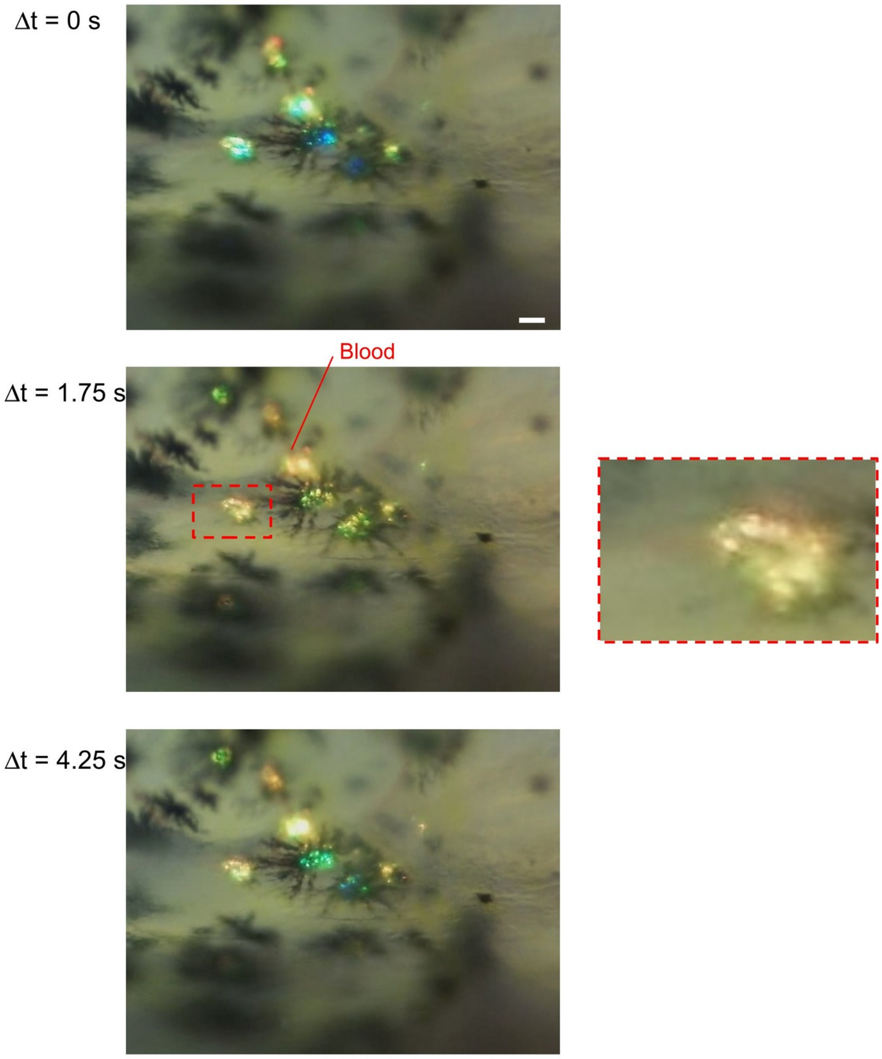


**FIGURE S19 |** An evidence of blood circulation getting close to IrH on xanthophore and YC. In about four seconds of the video playback, it was found that red path of blood appeared in two IrHs, as shown in middle picture. Right extraction image is a magnification of broken red box. Scale bar, 100 μm.

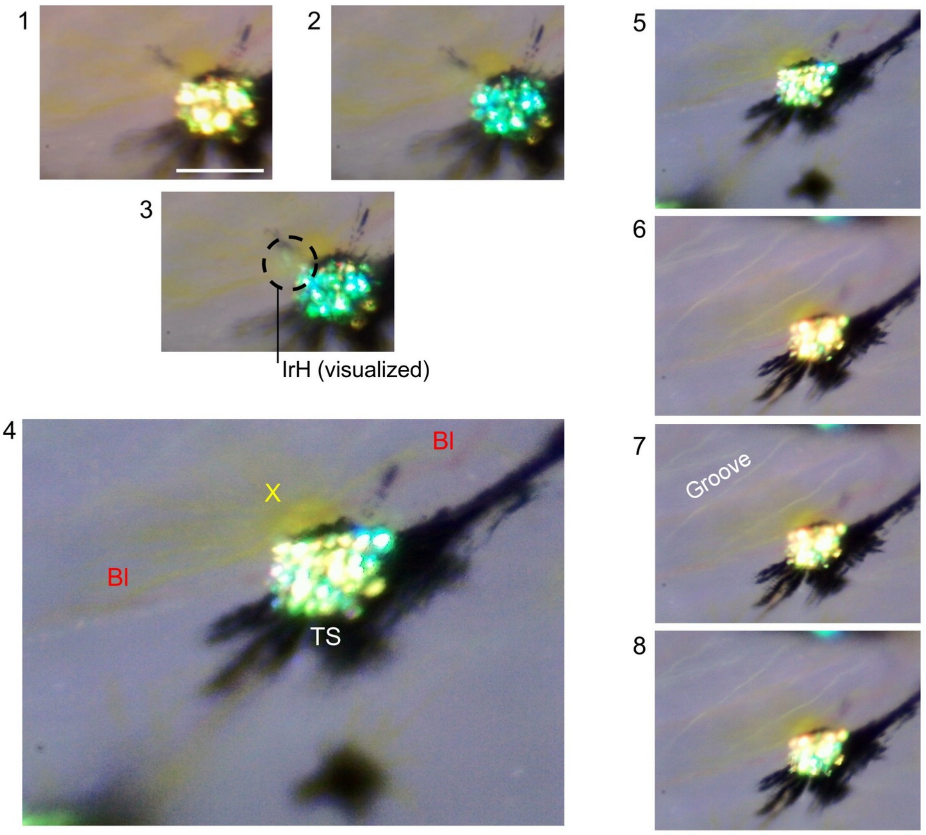


**FIGURE S20 |** Eight pictures showing micro-fluidic dynamics around a twinkling spot (TS). Pictures 1 and 6-8 shows the color of surrounding area of spots shifted to red when twinkling occurred. This supports the evidence that blood volume increment in the skin tissue enhances the twinkling intensity at TS. Picture 3 shows green stream appearing in the IrH layer (inside broken circle). In picture 4, a xanthophore is stretching along blood vessel (Bl). Most of white lines in pictures 5-8 are grooves of scale. Xanthophores stretching along the scale groove were frequently observed. Scale bar, 100 μm.

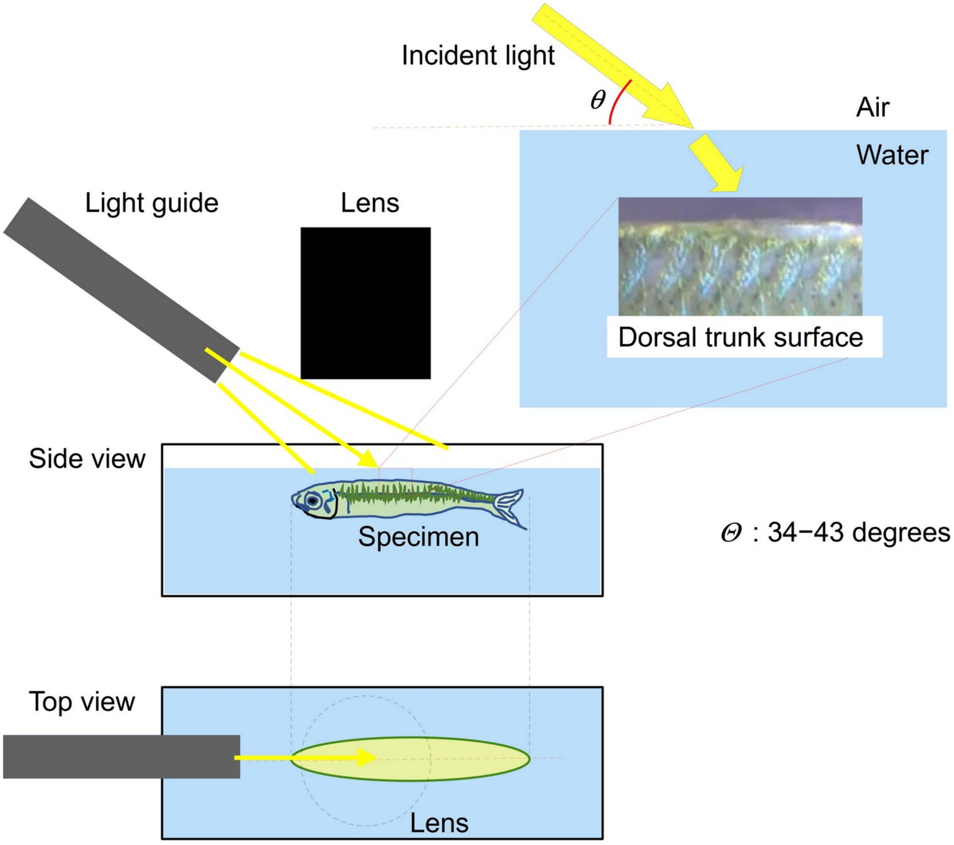


**FIGURE S21** | Schematic of the observation system for skin tissue of silverside fish under anesthesia control. Configurations of incident light and a specimen under water surface is drawn,

**Table S1. Overview of appearance of key functions on epidermis of sliverside fish belonging to a population of an experiment series.** A simple evaluation of appearance of the epidermal microcirculation functions was carried out on the population of fish captured between October 2020 to October 2024. Explanation of abbreviation used here is shown in the bottom of table.

| Species | Specimen number | **F-1** | **F-2** | **F-3** | **F-4** | **F-5** | **F-6** | **F-7** | Data presentation |
|---|---|---|---|---|---|---|---|---|---|
| *A. lacunosus* | spm20201007-1 | 〇 | 〇 | 〇 | NA | NA | NA | NA | * |
| *A. lacunosus* | spm20201007-2 | 〇 | 〇 | 〇 | NA | NA | NA | NA | Fig. 2d, Fig. S7 |
| *A. lacunosus* | spm20201007-3 | 〇 | 〇 | NA | NA | NA | NA | 〇 | * |
| *A. lacunosus* | spm20201007-4 | 〇 | 〇 | NA | NA | NA | NA | 〇 | * |
| *A. lacunosus* | spm20201007-5 | 〇 | 〇 | NA | NA | NA | NA | 〇 | * |
| *A. lacunosus* | spm20201007-6 | 〇 | 〇 | 〇 | NA | NA | NA | NA | * |
| *A. lacunosus* | spm20201211-1 | 〇 | 〇 | 〇 | - | - | 〇 | 〇 | Fig. S14 |
| *A. lacunosus* | spm20211106 | 〇 | 〇 | - | - | - | - | 〇 | Fig. S9 |
| *A. lacunosus* | spm20211108 | 〇 | 〇 | 〇 | 〇 | - | - | 〇 | Fig. 3d, 3e, Fig. S5 |
| *A. lacunosus* | spm20211109 | 〇 | 〇 | - | 〇 | - | - | 〇 | Fig. 1e |
| *A. lacunosus* | spm20211128 | 〇 | 〇 | 〇 | - | 〇 | 〇 | 〇 | Fig. 3c |
| *A. lacunosus* | spm20211129 | 〇 | NA | - | NA | NA | NA | 〇 | Fig. 1d, Fig. S3 |
| *H. valenciennei* | spm20230705-1 | 〇 | 〇 | - | - | - | - | 〇 | * |
| *H. valenciennei* | spm20230705-2 | 〇 | 〇 | - | - | - | - | 〇 | * |
| *H. valenciennei* | spm20230705-3 | 〇 | 〇 | - | - | 〇 | - | 〇 | * |
| *H. valenciennei* | spm20230705-4 | 〇 | 〇 | - | - | 〇 | - | 〇 | Fig. S12 |
| *H. valenciennei* | spm20230705-5 | 〇 | 〇 | 〇 | - | 〇 | - | 〇 | Fig. S4 |
| *H. tsurugae* | spm20230722-1 | 〇 | 〇 | 〇 | - | 〇 | 〇 | 〇 | Fig. S5c |
| *H. tsurugae* | spm20230722-2 | 〇 | 〇 | 〇 | 〇 | 〇 | 〇 | 〇 | * |
| *H. tsurugae* | spm20230722-3 | 〇 | 〇 | 〇 | - | - | 〇 | 〇 | * |
| *H. tsurugae* | spm20230722-4 | 〇 | 〇 | - | - | - | - | 〇 | * |
| *H. tsurugae* | spm20230722-5 | 〇 | 〇 | 〇 | - | 〇 | 〇 | 〇 | Fig. 1a |
| *H. tsurugae* | spm20230723-1 | 〇 | 〇 | 〇 | 〇 | 〇 | 〇 | 〇 | Fig. 3f, FigS2 |
| *H. tsurugae* | spm20230723-2 | 〇 | 〇 | - | - | 〇 | 〇 | 〇 | * |
| *H. tsurugae* | spm20230723-3 | 〇 | 〇 | 〇 | 〇 | 〇 | 〇 | 〇 | * |
| *H. tsurugae* | spm20230723-4 | 〇 | 〇 | 〇 | 〇 | 〇 | - | 〇 | * |
| *H. valenciennei* | spm20231004-1 | 〇 | 〇 | 〇 | 〇 | 〇 | 〇 | 〇 | Fig. 2a |
| *H. valenciennei* | spm20231004-2 | 〇 | 〇 | 〇 | - | 〇 | 〇 | 〇 | * |
| *H. valenciennei* | spm20231004-3 | 〇 | 〇 | 〇 | 〇 | 〇 | 〇 | 〇 | Fig. S1 |
| *H. valenciennei* | spm20231126-1 | 〇 | 〇 | 〇 | 〇 | - | 〇 | 〇 | Fig. S18, Fig. 2h, Fig. S20. Fig. S10, Fig. S15 |
| *H. valenciennei* | spm20231127-1 | 〇 | 〇 | 〇 | 〇 | 〇 | 〇 | 〇 | Fig. S17a |
| *H. valenciennei* | spm20231127-2 | 〇 | 〇 | 〇 | - | 〇 | - | 〇 | Fig. S17b |
| *H. tsurugae* | spm20240529-1 | 〇 | 〇 | 〇 | - | - | 〇 | 〇 | * |
| *H. tsurugae* | spm20240529-2 | 〇 | 〇 | 〇 | - | - | 〇 | 〇 | * |
| *H. tsurugae* | spm20240529-3 | 〇 | 〇 | 〇 | - | 〇 | - | 〇 | * |
| *H. valenciennei* | spm20240529-4 | 〇 | 〇 | 〇 | - | 〇 | 〇 | 〇 | * |
| *H. tsurugae* | spm20240529-5 | 〇 | 〇 | 〇 | - | - | 〇 | 〇 | * |
| *H. tsurugae* | spm20240529-6 | 〇 | 〇 | 〇 | - | 〇 | 〇 | 〇 | * |
| *H. tsurugae* | spm20240529-7 | 〇 | 〇 | 〇 | - | 〇 | - | 〇 | * |

| Species | Specimen number | **F-1** | **F-2** | **F-3** | **F-4** | **F-5** | **F-6** | **F-7** | Data presentation |
|---|---|---|---|---|---|---|---|---|---|
| *H. tsurugae* | spm20240529-8 | ○ | ○ | ○ | ○ | ○ | ○ | ○ | * |
| *H. tsurugae* | spm20240529-9 | ○ | ○ | ○ | - | ○ | - | ○ | Fig. 1 b, c |
| *H. tsurugae* | spm20240529-10 | ○ | ○ | ○ | ○ | ○ | ○ | ○ | * |
| *H. tsurugae* | spm20240530-1 | ○ | ○ | ○ | - | ○ | - | ○ | * |
| *H. tsurugae* | spm20240530-2 | ○ | ○ | - | - | ○ | ○ | ○ | Fig. S13 |
| *H. tsurugae* | spm20240530-3 | ○ | ○ | ○ | - | ○ | ○ | ○ | * |
| *H. tsurugae* | spm20240530-4 | ○ | ○ | - | - | ○ | ○ | ○ | * |
| *H. valenciennei* | spm20240925-1 | ○ | ○ | ○ | - | - | ○ | ○ | Fig. 3h |
| *H. valenciennei* | spm20240925-2 | ○ | ○ | ○ | - | - | - | ○ | * |
| *H. valenciennei* | spm20240925-3 | ○ | ○ | ○ | - | - | - | ○ | * |
| *H. valenciennei* | spm20240925-4 | ○ | ○ | ○ | ○ | - | - | ○ | * |
| *H. valenciennei* | spm20240925-5 | ○ | - | ○ | - | - | - | ○ | * |
| *H. valenciennei* | spm20240925-6 | ○ | ○ | ○ | - | ○ | ○ | ○ | * |
| *H. valenciennei* | spm20240925-7 | ○ | ○ | - | - | - | - | ○ | * |
| *H. valenciennei* | spm20240925-8 | ○ | ○ | ○ | ○ | - | ○ | ○ | Fig. S8, S11 |
| *H. valenciennei* | spm20240928-1 | ○ | ○ | - | - | - | - | ○ | Fig. S4, S6, S19 |
| *H. valenciennei* | spm20240928-2 | ○ | ○ | ○ | - | - | - | ○ | * |
| *H. valenciennei* | spm20240928-3 | ○ | ○ | ○ | - | - | - | ○ | * |
| *H. valenciennei* | spm20240928-4 | ○ | ○ | - | - | - | - | ○ | * |
| *H. valenciennei* | spm20240928-5 | ○ | ○ | ○ | - | - | - | ○ | Fig. S6 |
| *H. valenciennei* | spm20240928-6 | ○ | ○ | ○ | - | ○ | ○ | ○ | * |
| *H. valenciennei* | spm20241011-1 | ○ | ○ | ○ | - | - | ○ | - | Fig. 3g |
| *H. valenciennei* | spm20241011-2 | ○ | ○ | - | - | - | - | ○ | * |
| *H. valenciennei* | spm20241011-3 | ○ | ○ | - | - | - | - | ○ | * |
| *H. valenciennei* | spm20241011-4 | ○ | ○ | - | - | - | - | ○ | * |
| *H. valenciennei* | spm20241011-5 | ○ | - | - | - | - | - | ○ | * |
| *H. valenciennei* | spm20241011-6 | ○ | - | - | - | - | - | ○ | * |
| *H. valenciennei* | spm20241011-7 | ○ | - | ○ | - | - | - | ○ | * |
| *H. valenciennei* | spm241017-mag | ○ | ○ | ○ | - | - | - | NA | Fig. S16 |
| *A. lacunosus* | spm2020XX | ○ | NA | NA | NA | NA | NA | ○ | Fig. 2e |
| *H. tsurugae* | Spm202324XX-1 | ○ | NA | NA | NA | NA | NA | ○ | Fig. 2c |
| *H. tsurugae* | Spm202324XX-2 | ○ | ○ | NA | NA | NA | NA | NA | Fig. 2f |
| *H. tsurugae* | Spm202324XX-3 | ○ | ○ | ○ | NA | NA | NA | NA | Fig. 2i |

Functions

**F-1** **Capillaries (white or yellow) exist**
**F-2** **Twinkling spots**
**F-3** **Brilliant yellow capillary exists**
**F-4** **Grid pattern appears on a twinkling spot**
**F-5** **Twinkling on IrH in the absence of melanophore**
**F-6** **Fluid pressure propagation pattern**
**F-7** **IrH membrane exists**
○ Function was detected in specimen
NA Not available due to observation position of tissue and illumination direction
- Function can not be identified in specimen
* Detailed data are not shown in this manuscript

**Table S2. Reproducibility of the function in the population of specimens for this study.** Frequency of the appearance of function shown in Table S1. is converted to the reproducibility excluding the number of NA in each function.

Reproducibility (%)

| Capillaries (white or yellow) exist | Twinkling spots | Brilliant yellow capillary exists | Grid pattern appears on a twinkling spot | Twinkling on IrH in the absence of melanophore | Fluid pressure propagation pattern | IrH membrane exists |
|---|---|---|---|---|---|---|
| 100 | 94.2 | 71.2 | 22.9 | 47.5 | 47.5 | 98.5 |

Tables S1 and S2 show overview of the reproducibility of the appearance of the focused functions which were discovered in three species of silverside fish. Existence of the discovered functions of chromatophores and epidermal microcirculation system that was referred in the main text were evaluated for individual specimen. There were other captured samples than the specimens providing the figure presentation in this paper. The same kind of observation experiment was carried out on the population of specimens captured in the same day. The evaluation of reproducibility of the functions was also carried out on the same day specimens whose detailed data were not presented as figures in the paper (main text and supporting information). All specimens belong to the dataset groups were evaluated in the same manner with the data-presented specimens.
